# Phosphorylation-driven Targeted Protein Degradation of Oncogenic β-catenin

**DOI:** 10.64898/2026.03.16.712096

**Authors:** Lucie Wolf, Juline Poirson, Graham MacLeod, Sichun Lin, Yun Hye Kim, Maira P Almeida, Mikko Taipale, Stéphane Angers

## Abstract

Therapeutic strategies to inhibit the Wnt signalling pathway for cancer treatment have, so far, failed to advance to the clinic. Induced-proximity drugs are revolutionizing our ability to tackle targets previously considered undruggable. Here, we used an unbiased genome-scale approach to identify induced-proximity protein candidates that inhibit the central Wnt signalling effector β-catenin in colorectal cancer cells. While the identification of several E3 ubiquitin ligases validated our approach, we uncovered that inducing proximity to members of the Casein kinase I (CSNK1) family leads to β-catenin degradation and inhibits the growth of colorectal cancer cells harbouring Wnt pathway mutations. We show that β-catenin degradation induced by CSNK1 proximity is kinase activity- and proteasome-dependent. We propose that the formation of a neo-degron, through kinase recruitment, can expand induced-proximity drug targeting strategies.

## INTRODUCTION

The β-catenin-dependent Wnt signalling pathway regulates stem and progenitor cell activity during embryonic development and maintains tissue homeostasis in adults (Rim et al., 2022; Wolf and Boutros, 2023; Maurice and Angers, 2025). Binding of Wnt ligands to their receptors at the cell surface activates an intracellular signalling pathway that culminates in the stabilisation and accumulation of the central effector protein β-catenin in the cytoplasm and the nucleus. This induces transcriptional regulation of downstream signalling programmes and regulates proliferation, cell survival and other cellular effects in a context-dependent manner (Maurice and Angers, 2025; Söderholm and Cantù, 2021). In the absence of Wnt ligands, when the signalling pathway is inactive, cytoplasmic β-catenin protein abundance is tightly regulated through post-translational modifications by the so-called ‘destruction complex’, which is formed by the scaffolding proteins Adenomatous Polyposis Coli (APC) and AXIN, as well as the kinases Glycogen Synthase Kinase 3 α and β (GSK3A/B) and Casein kinase Iα (CSNK1A1). This protein complex catalyses the phosphorylation of β-catenin to earmark it for ubiquitination and subsequent degradation by the proteasome (Maurice and Angers, 2025; van Kappel and Maurice, 2017). Mutations affecting components of the destruction complex or β-catenin itself, which results in ligand-independent β-catenin stabilisation, are frequently detected in cancer, most notably the neoplasms affecting the gastrointestinal tract (Parsons et al., 2021; Werner et al., 2023; Zhan et al., 2017). In fact, these mutations occur in nearly all cases of colorectal cancer and promote cell survival and proliferation of progenitor cells (Schatoff et al., 2017; Söderholm and Cantù, 2021; Yaeger et al., 2018). Importantly, studies in mouse models of colon cancer exhibiting high Wnt/β-catenin signalling showed that lowering β-catenin levels in these tumours prevents cancer progression and restores homeostasis (Dow et al., 2015; Scholer-Dahirel et al., 2011). Tumours therefore rely on continuing β-catenin signalling for growth, suggesting that inhibition of Wnt/β-catenin signalling to block cancer progression is a logical therapeutic target, but, to date, no drugs have been clinically approved (Maurice and Angers, 2025; Parsons et al., 2021; Wang et al., 2021). This shortcoming is attributed to a wide variety of unwanted on-target effects stemming from systemic Wnt signalling modulation and its impact on tissue stem cell activity, for example affecting bone and intestinal homeostasis (Liu et al., 2022; Maurice and Angers, 2025; Zhan et al., 2017). More fundamentally, this difficulty is also associated with targeting downstream Wnt pathway components that function mostly through protein-protein interactions that are harder to target with small molecules than enzymes.

Targeted Protein Degradation (TPD) based on bifunctional molecules developed to promote the proximity of a protein of interest with components of degradation pathways such as the ubiquitin system is now a proven strategy to tackle, previously considered, undruggable targets. The most prominent examples are PROteolysis-TArgeting Chimeras (PROTACs), and molecular glues, which both link target proteins to E3 ubiquitin ligases, in such a way to promote their ubiquitination and proteasomal degradation (Békés et al., 2022; Li and Crews, 2022; Schreiber, 2021; Zhao et al., 2022). While PROTACs and molecular glues have been designed to successfully target a wealth of targets, with promising molecules advancing in clinical trials, the general concept is still blossoming (Chirnomas et al., 2023; Hinterndorfer et al., 2025; Jan et al., 2021; King et al., 2025; Mayor-Ruiz and Winter, 2019). For instance, PROTACs and molecular glues only use a limited number of the over 600 E3 ubiquitin ligases available in the human genome, and mostly rely on the well characterised E3s VHL and CRBN for which chemical warheads exist (Békés et al., 2022; Bricelj et al., 2021; Chirnomas et al., 2023; King et al., 2025). Beyond ubiquitin ligases, inducing the proximity with other classes of proteins has also been leveraged to regulate the activity of targets of interest. For instance, researchers were able to modulate cellular signalling pathways through synthetic tyrosine phosphorylation (Pergu et al., 2023; Shoba et al., 2022; Siriwardena et al., 2020) or to induce apoptosis by repurposing transcriptional kinases (Sarott et al., 2024).

Here, we leveraged a recently developed induced-proximity screening strategy (Poirson et al., 2024) to identify proteins from a near-proteome-wide library of around 15,000 Open Reading Frames (ORFs) that would regulate β-catenin abundance and function when brought into its proximity. We performed this screen in human colon cancer cells where β-catenin signalling is essential for their growth. Our results expectedly revealed many E3 ubiquitin ligases and known TPD effectors, validating our approach. To our surprise, we found that members of the CSNK1 family were amongst the strongest regulators of β-catenin protein levels. We confirmed that the CSNK1-dependent degradation mechanism is proximity-, kinase-activity- and Ubiquitin-Proteasome System (UPS)-dependent. We propose that the rescue of the β-catenin phospho-degron, which is inhibited in colorectal cancers, through the forced recruitment of a kinase constitutes a novel TPD strategy that we predict could be expanded to other targets and disease indications.

## RESULTS

### Induced-proximity screens identify potent degraders of β-catenin in colorectal cancer cells

To identify proteins within the human proteome that promote β-catenin degradation when brought into close proximity, we used a recently described induced-proximity screening system (Poirson et al., 2024). In brief, a near-proteome-wide library of 14,821 human Open Reading Frames (ORFs) covering all protein classes and functions were expressed as fusion proteins with an anti-GFP nanobody (vhhGFP) at the C-terminus. VhhGFP strongly binds GFP and related fluorophores and in this way constitutively recruits proteins encoded by each ORF in the library into close proximity to any GFP-tagged protein (**Fig 1A**). To use this system, we used gene editing to tag both alleles of *CTNNB1* (the gene encoding β-catenin) in the human colon cancer cell line DLD-1 with GFP variants. This cell line carries inactivating *APC* mutations, a component of the destruction complex, which leads to ligand-independent constitutive stabilisation of β-catenin and hyperactivation of the Wnt signalling pathway. Using the tagging strategy previously described by de Man et al. (de Man et al., 2021, **Fig 1B**), we tagged β-catenin at the N-terminus just upstream of its start codon in a stepwise process: one allele was first tagged with enhanced Green Fluorescent Protein (eGFP) and the second allele was subsequently tagged with enhanced Blue Fluorescent Protein 2 (eBFP2). The nucleotide sequences of these two fluorophores differ only in 22 bp, corresponding to 14 amino acids, and the vhhGFP binds to both. Thus, we were able to investigate regulation of both eGFP- and eBFP2-tagged β-catenin using the ORFeome-vhhGFP library.

**Fig. 1:**
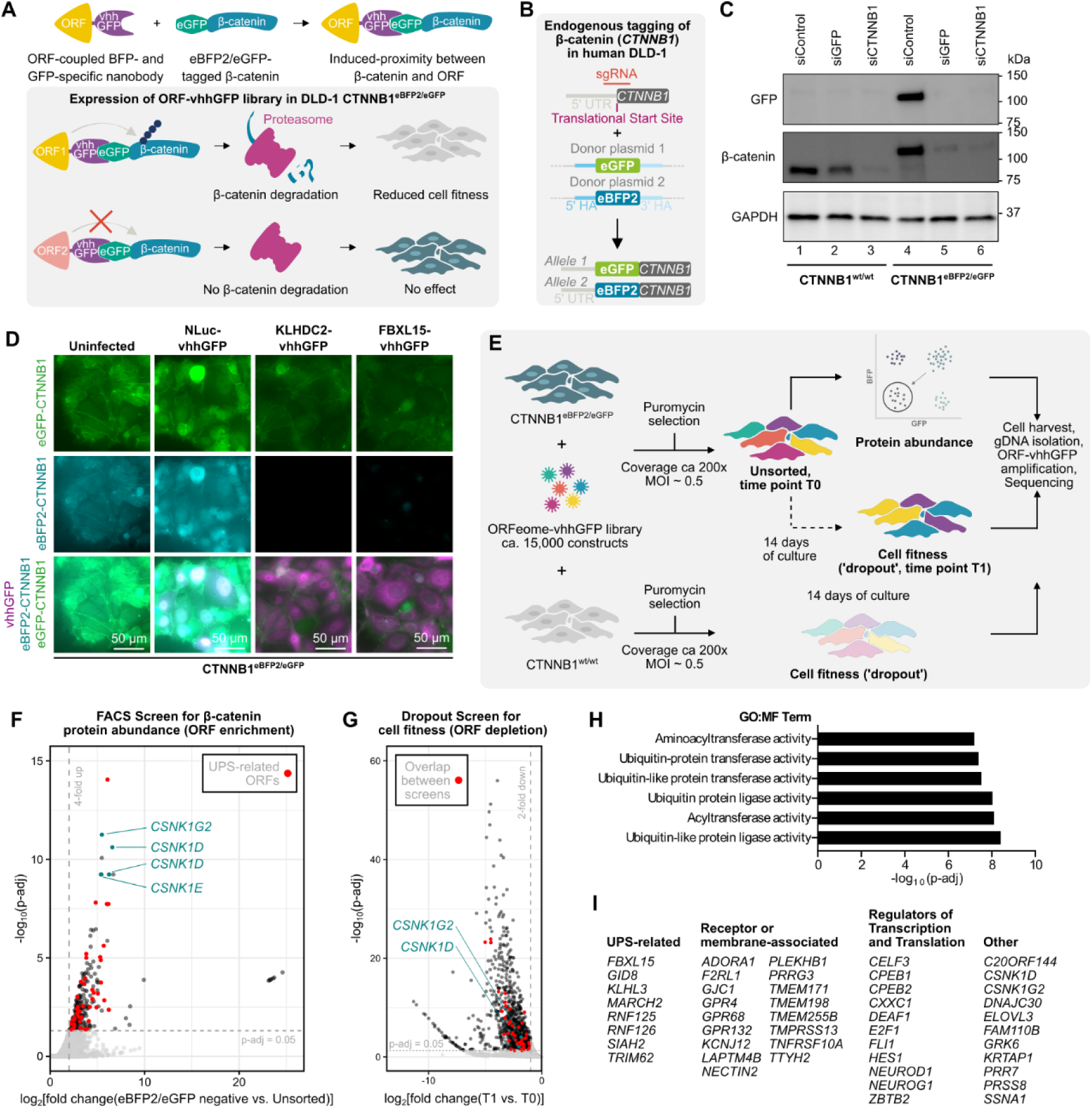
Induced proximity screening strategy identifies potent regulators of β-catenin for targeted protein degradation. **A)** Schematic depiction of possible outcomes resulting from induced proximity between the anti-GFP nanobody (vhhGFP)-coupled Open Reading Frame (ORF) library and endogenously double tagged eBFP2/eGFP-β-catenin. vhhGFP binds both eGFP and eBFP2. **B)** The colorectal cancer cell line DLD-1 was endogenously tagged with eGFP and eBFP2 upstream of the β-catenin (*CTNNB1* gene) start codon using CRISPR/Cas9. sgRNA – single guide RNA, UTR – untranslated region, HA – homology arm **C)** β-catenin protein levels are reduced after siGFP transfection in eBFP2/eGFP-β-catenin expressing DLD-1 cells. Untagged or double tagged DLD-1 CTNNB1^eBFP2/eGFP^ cells (clone #28) were transfected with the indicated siRNA and harvested for Western Blot analysis 72 h later. siGFP targeted both eGFP and eBFP2 due to sequence similarities between the two fluorophores. Representative Western Blot of n = 3 biological replicates. kDa = Kilodalton, GAPDH served as loading control. **D)** eBFP2/eGFP-β-catenin is degraded by induced proximity to E3 ubiquitin ligase adapters fused with vhhGFP. Immunofluorescence imaging of endogenous eBFP2- and eGFP-β-catenin in the presence of NanoLuc luciferse (NLuc)-, KLHDC2-, or FBXL15-vhhGFP. **E)** Screening strategy for the Fluorescence Activated Cell Sorting (FACS) β-catenin protein abundance screen, the cell fitness (dropout) screen and the control cell fitness (dropout) screen in untagged cells. MOI – multiplicity of infection. **F, G)** Volcano plots depicting significant hits from the FACS and dropout screens. 47 hits were found in both screens **(I)**. The FACS screen had a significant enrichment for Ubiquitin-related Molecular Function (MF) Gene Ontology (GO) terms (**H).** UPS - Ubiquitin-Proteasome System

Sanger sequencing confirmed seamless integration of both eGFP and eBFP2 at the intended genomic loci in three independent single cell clones (**Suppl Fig 1**). Extensive validation of these clones confirmed that the expression (**Fig 1C, Suppl Fig 2A**), localisation (**Fig 1D**) and function of the tagged β-catenin alleles were indistinguishable from wild-type cells (**Suppl Fig 2B, Suppl Fig 2C).** β-catenin is a fitness gene required for the proliferation of most colorectal cancer cell lines (The DepMap Portal, www.depmap.org). The double tagged eGFP/eBFP2-β-catenin DLD-1 cells still rely on β-catenin for their proliferation as knock-out of *CTNNB1* led to proliferation defects and mutant cells dropped out from the population when competing with unedited control cells (**Suppl Fig 2D**).

As proof of concept that β-catenin protein levels can be regulated through induced-proximity in these double-tagged cells, we overexpressed the E3 ubiquitin ligase substrate adapters KLHDC2 and FBXL15 fused to vhhGFP and investigated β-catenin protein abundance using fluorescence microscopy. KLHDC2 was previously reported as a strong induced-proximity degrader of β-catenin (Röth et al., 2023) and FBXL15 was one of the strongest degraders described by Poirson et al. (Poirson et al., 2024). As expected, both eGFP- and eBFP2-tagged β-catenin were markedly reduced in cells that stained positive for the E3 ligase adaptors (**Fig 1D**).

Using the double-tagged eGFP/eBFP2-β-catenin DLD-1 cells in combination with the ORFeome-vhhGFP library, we then conducted two independent screens to identify potent regulators of β-catenin protein levels and function. The first was a FACS-based phenotypic screen where eGFP/eBFP2 expression served as a proxy for β-catenin protein abundance, and the second was a cell fitness (dropout) screen since disruption of β-catenin function inhibits DLD-1 cell proliferation (**Suppl Fig 2B**). We also carried a control cell fitness screen in wild-type DLD-1 cells to exclude ORFs that affected DLD-1 proliferation independent of their induced proximity to β-catenin (**Fig 1E**).

For both screens, we infected eGFP/eBFP2-β-catenin DLD-1 cells with lentivirus expressing the ORFeome-vhhGFP library so that statistically each cell only expressed one ORF-vhhGFP fusion protein (Multiplicity of Infection, MOI, 0.5). For the protein abundance screen, we selected the infected cells with puromycin, expanded the surviving cells for three more days and used FACS to collect cells that had lost both eBFP2- and eGFP-expression (the double negative population), assuming these were the cells were induced proximity to the respective ORF had led to β-catenin degradation. To increase stringency, we focused on cells where both alleles of β-catenin were degraded (loss of both eBFP2 and eGFP) and excluded the few cells where only one fluorophore was affected. Amongst the 255 statistically significant hits identified in the protein abundance screen, 46 are associated with the Ubiquitin Proteasome System and Gene Ontology (GO) term analysis confirmed Ubiquitin-related Molecular Function (MF) amongst the highest statistically significantly enriched terms (**Fig 1F,H**).

For the fitness screen, cells were cultured for 14 days after puromycin selection and sgRNA abundance was quantified at T0 (baseline) and at endpoint (T1) using Next Generation Sequencing. 862 statistically significant hits were found to affect cell fitness selectively in the eGFP/eBFP2-β-catenin DLD-1 cells when compared to untagged cells. Many of these hits were histone, chromatin or transcription factor related, suggesting a potential impact on chromatin when these ORFs are forced into proximity with β-catenin. Importantly, the two screens had 47 hits in common, of which 8 were UPS related, 17 were membrane or membrane-associated proteins, 11 were involved in transcription or translation and 11 associated with other miscellaneous functions (**Fig 1G,I**). Interestingly, several hits, including GID8 and members of the FBXL and KLHL E3 ubiquitin ligase families, were previously found in other induced proximity screens, indicating strong, universal degraders that are likely not specific to a given target (Poirson et al., 2024). However, several high-ranking candidates appear to be unique and robust induced-proximity-dependent degrader specific to β-catenin. Given that the E3 ubiquitin ligase SIAH2 and members of the Casein-kinase I family (CSNK1) had been previously linked to Wnt/β-catenin signalling we set out to further characterize these candidate hits (Amit et al., 2002; Liu et al., 2002; O’Connell et al., 2013; Topol et al., 2003; van Kappel and Maurice, 2017).

### Induced proximity to the kinase CSNK1D or the E3 ubiquitin ligase SIAH2 degrades β-catenin

To validate the screen results we infected eGFP/eBFP2-β-catenin DLD-1 cells with the individual ORF-vhhGFP fusions or, as controls, the same ORFs fused to the NbALFA nanobody, which is specific for the ALFA-tag and does not target eGFP, eBFP2, or β-catenin (Götzke et al., 2019) and determined the impact on β-catenin abundance using flow cytometry (**Fig 2A**). As controls, we included uninfected cells and cells expressing Renilla Luciferase (RLuc)- or NanoLuciferase (NLuc)-vhhGFP fusion proteins. While RLuc-vhhGFP expression did not lead to a significant change in β-catenin levels when compared to RLuc-NbALFA, expression of NLuc-vhhGFP led to a small but detectable decrease in β-catenin abundance. This is consistent with vhhGFP binding stabilising GFP-tagged proteins (Poirson et al., 2024). Expression of CSNK1D and SIAH2 tagged with vhhGFP led to robust decrease in β-catenin to levels surpassing the FBXL15 positive control (**Fig 2B**). Recruitment of VHL, which is often used for PROTACs (Békés et al., 2022), only had a minor effect similar to the NLuc-vhhGFP control (**Fig 2B**). Further validating these results, we monitored β-catenin levels using Western blotting and detected robust degradation of β-catenin by CSNK1D- and SIAH2-vhhGFP but no degradation by the NbALFA fusion proteins (**Fig 2C**). The effect of FBXL15- and VHL-vhhGFP were less significant using this assay. We confirmed that β-catenin degradation was post-translationally regulated since expression of CSNK1D-and SIAH2-vhhGFP had no effect on *CTNNB1* (β-catenin) mRNA levels (**Fig 2D**). Consistent with on-target and sustained degradation of β-catenin, CSNK1D- and SIAH2-vhhGFP expression led to robust inhibition of the universal β-catenin target gene *AXIN2* (**Fig 2D**). Since β-catenin is essential for the growth of colorectal cancer cells, we next designed an experiment to evaluate the impact of ORF-vhhGFP expression on the growth fitness of eGFP/eBFP2-β-catenin DLD-1 cells. For this, we infected eGFP/eBFP2-β-catenin DLD-1 cells with virus directing the co-expression of CSNK1D- or SIAH2-vhhGFP together with mNeonGreen (a very bright green fluorophore) or with NLuc-vhhGFP and TagRFP (a very bright red fluorophore) from bicistronic constructs. Importantly, vhhGFP does not interact with these fluorophores and they are much brighter than the endogenously tagged eGFP/eBFP2-β-catenin. Following selection with puromycin, green CSNK1D-vhhGFP or SIAH2-vhhGFP expressing cells were mixed with control red cells expressing NLuc-vhhGFP and their proliferation analysed using the Incucyte automatic time-lapse fluorescent microscope, which only detected the fiducial markers TagRFP and mNeonGreen. In these co-culture experiments, the green area corresponding to CSNK1D-vhhGFP or SIAH2-vhhGFP expressing cells barely increased over the course of 6 days, indicating severely impaired proliferation compared to the control red cells expressing NLuc-vhhGFP, which steadily increased their occupied area (**Fig 2E**). For orthogonal validation in an alternative dimerisation system without eGFP/eBFP2, we created a stable HEK293 Flp-In T-REx cell line expressing HiBiT-ALP5-β-catenin. NbALFA binds the Alp5 tag (Götzke et al., 2019; Kilisch et al., 2021) and expression of CSNK1D- and SIAH2-NbALFA fusion effectors significantly reduced HiBiT-ALP5-β-catenin protein levels compared to NLuc-NbALFA in transfected cells, confirming our results in DLD-1 cells (**Suppl Fig S3**). We conclude that induced-proximity to CSNK1D and SIAH2 potently regulated β-catenin protein abundance, target gene expression and the proliferation of colorectal cancer cells.

**Fig. 2:**
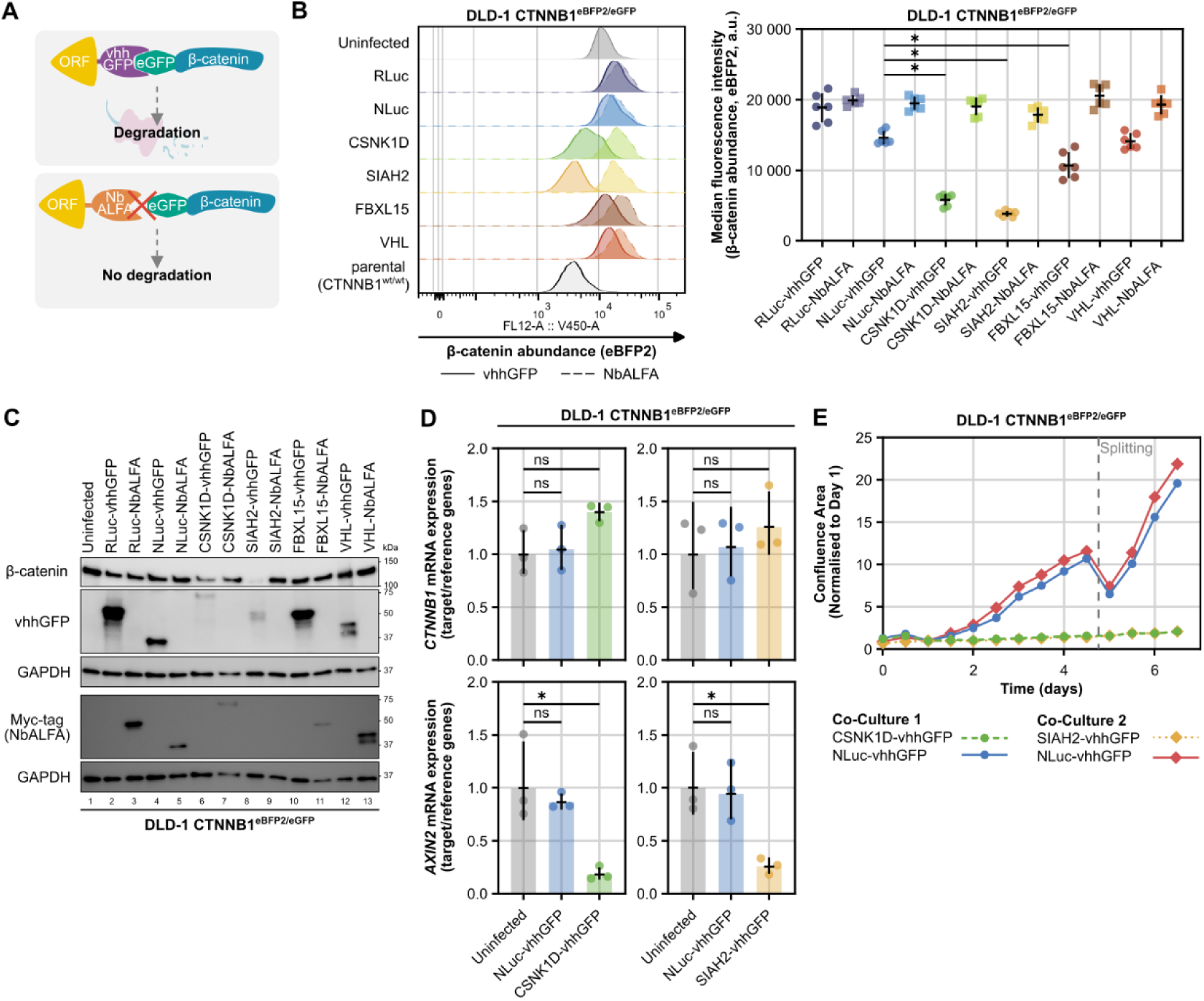
The kinase CSNK1D and the E3 ubiquitin ligase SIAH2 are strong regulators of β-catenin. **A)** Schematic depiction of how eBFP2/eGFP-β-catenin is brought into close proximity to anti-GFP nanobody (vhhGFP)-coupled Open Reading Frames (ORF) but not to anti-ALFA-tag nanobody (NbALFA)-coupled ORFs. vhhGFP binds both eGFP and eBFP2. **B, C, D)** Hits from the induced-proximity screens degrade β-catenin protein and regulate β-catenin target gene expression through post-translational mechanisms. DLD-1 CTNNB1^eBFP2/eGFP^ cells (clone #28) were infected with virus expressing the indicated -vhhGFP or -NbALFA fusion proteins, selected with puromycin for 2 days and then analysed. **B)** eBFP2 expression was measured by flow cytometry as read-out for β-catenin abundance. On the left, histograms from one experiment of clone #28 are shown. On the right, individual data points from two independent experiments with 3 clones (#28, #83, #92) are shown with mean and standard deviation. A minimum of 10 000 cells per condition were recorded in the final gate. Statistical significance was assessed by one-way ANOVA followed by Dunnett’s multiple comparisons test comparing all -vhhGFP groups to NLuc-vhhGFP. * Padj ≤ 0.0001, ns = not significant. **C)** Representative Western Blot of n = 3 biological replicates. kDa = Kilodalton, GAPDH served as loading control. **D)** *CTNNB1* and *AXIN2* mRNA expression was determined using reverse-transcription quantitative polymerase chain reaction (RT-qPCR). Fold change data was normalised to uninfected cells and *GAPDH*, *B2M* and *PPIA* served as reference genes. Individual data points from three independent experiments are shown with geometric mean and geometric standard deviation. Statistical significance was assessed by one-way ANOVA followed by Dunnett’s multiple comparisons test compared to uninfected cells. * Padj ≤ 0.05, ns = not significant. **E)** DLD-1 CTNNB1^eBFP2/eGFP^ cells (clone #28) were infected with virus expressing the indicated -vhhGFP fusion proteins and red (NLuc-vhhGFP, TagRFP) or green (CSNK1D-vhhGFP or SIAH2-vhhGFP, mNeonGreen) fluorophores. Then, NLuc-vhhGFP (TagRFP)- and CSNK1D-vhhGFP or SIAH2-vhhGFP (mNeonGreen)-expressing cells were mixed and imaged using a time-lapse microscope (Incucyte) over the course of 6 days to assess cell confluence. In this co-culture experiment, cells with strong regulation of β-catenin abundance (e.g. CSNK1D-vhhGFP or SIAH2-vhHGFP expressing cells) proliferated less than their control counterparts. Representative of n = 3 biological replicates. RLuc - Renilla luciferase; NLuc - NanoLuc luciferase; vhhGFP - nanobody binding to eGFP and eBFP2; NbALFA - nanobody recognising ALFA-tag; a.u. - arbitrary unit

### Small molecule-induced proximity between β-catenin and CSNK1D regulates Wnt signalling and inhibits colon cancer cell proliferation

The results so far indicating CSNK1D- or SIAH2-dependent degradation of β-catenin relied on the constitutive recruitment of ORF-vhhGFP fusions to eGFP/eBFP2-β-catenin (nanobody-antigen binding). We next established inducible recruitment systems using small molecules as proof-of-principle that a PROTAC strategy could be effective in this context. We first leveraged the small molecule-dependent dimerization of the FKBP12^F36V^ and FRB* domains mediated by Rapamycin or the Rapalog A/C Heterodimeriser (previously known as AP21967) (Bayle et al., 2006; Inoue et al., 2005; Muthuswamy et al., 1999). We employed the same endogenous tagging strategy as above to engineer a DLD-1 cell line in which both *CTNNB1* alleles are tagged at the N-terminus with eGFP-FRB* (**Fig 3A**). Upon expression of CSNK1D-FKBP12^F36V^, treatment of eGFP-FRB*-β-catenin DLD-1 cells with Rapamycin or the Rapalog A/C Heterodimeriser led to robust degradation of β-catenin whereas SIAH2-FKBP12^F36V^ expression only had mild effects in this system (**Fig 3B,C**). Strikingly, CSNK1D-FKBP12^F36V^ -dependent degradation of eGFP-FRB*-β-catenin induced by Rapamycin or the A/C Heterodimeriser impaired cell proliferation in long-term assays of up to 14 days (**Fig 3D**). We also used an orthogonal dimeriser system that leverages the inducible binding of FKBP12^F36V^ and HaloTag by the bivalent molecule PhosTAC7 (Chen et al., 2021). To do this, we endogenously tagged both *CTNNB1* alleles with HaloTag and ectopically expressed CSNK1D- or SIAH2-FKBP12^F36V^. PhosTAC7 treatment led to significant decrease in HaloTag-β-catenin levels when compared to controls (**Suppl Fig 4A,B,C**).

**Fig. 3:**
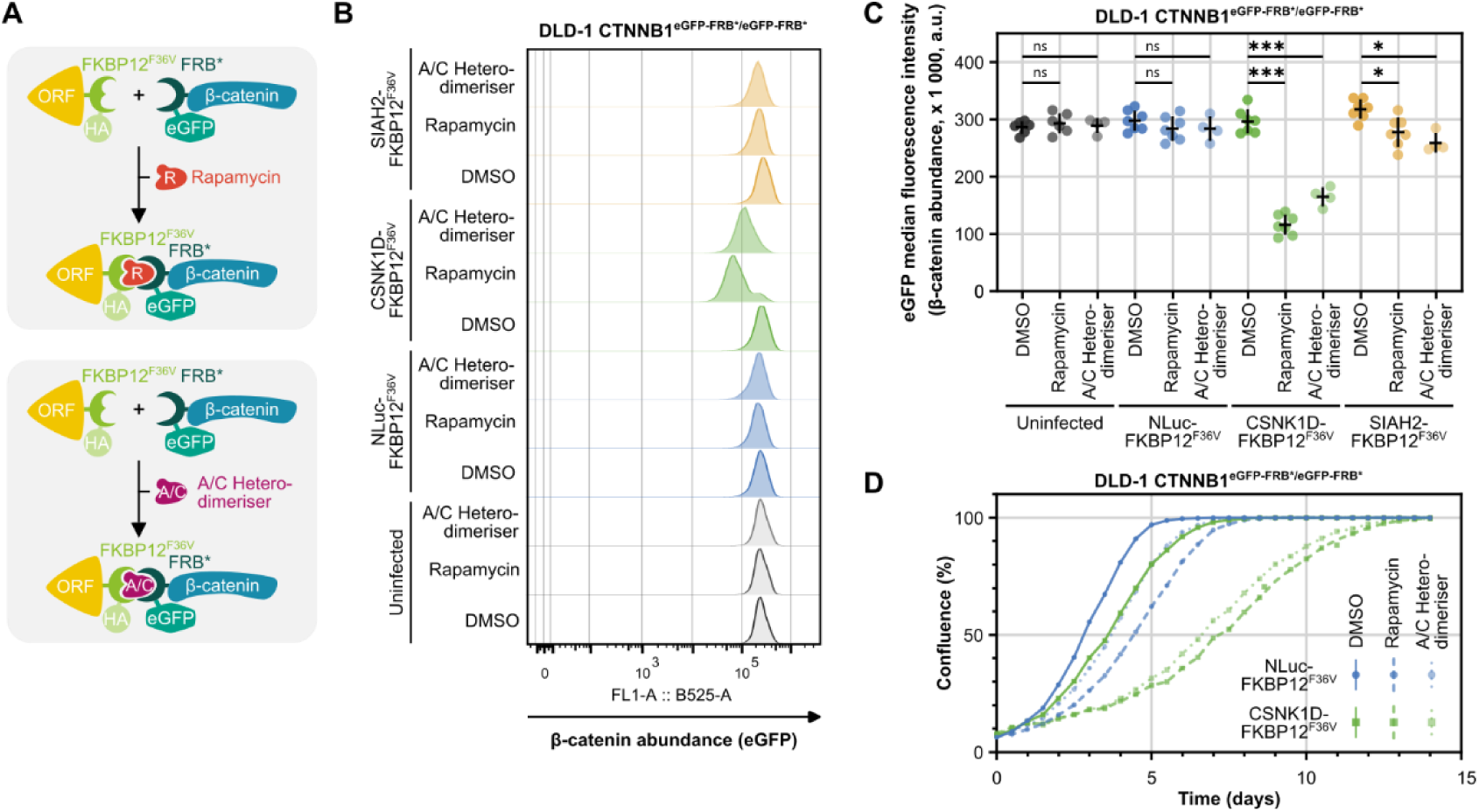
Chemically induced proximity to CSNK1D degrades β-catenin and inhibits growth in colorectal cancer cells. **A)** Schematic depiction of induced proximity between eGFP-FRB*-β-catenin and HA- and FKBP12^F36V^- tagged Open Reading Frames (ORF) using the small molecules Rapamycin or the Rapalog A/C Heterodimeriser. **B, C)** Rapamycin or A/C Heterodimeriser induced dimerisation of eGFP-FRB*-β-catenin and CSNK1D- or SIAH2-FKBP12^F36V^ leads to β-catenin degradation in DLD-1 colorectal cancer cells. Double tagged DLD-1 CTNNB1^eGFP-FRB*/eGFP-FRB*^ cells (clone #43) were infected with virus expressing the indicated -FKBP12^F36V^-HA fusion proteins, selected with puromycin for 2 days and then treated with Rapamycin (0.5 µM), A/C Heterodimeriser (1 µM) or equivalent volumes of DMSO for 24 h. Then, eGFP-FRB*-β-catenin abundance was quantified by flow cytometry. **B)** Histograms of eGFP intensity of one representative biological replicate are shown normalised to mode. **C)** Median fluorescence intensity of eGFP in n = 4 or 7 biological replicates with mean and standard deviation. A minimum of 8,800 cells per condition were recorded in the final gate. Statistical significance was assessed by one-way ANOVA followed by Dunnett’s multiple comparisons test comparing treatment groups to DMSO. * Padj ≤ 0.05, *** Padj ≤ 0.0001, ns = not significant. **D)** Double tagged DLD-1 CTNNB1^eGFP-FRB*/eGFP-FRB*^ cells (clone #43) expressing CSNK1D-FKBP12^F36V^ exhibit a strong growth defect when treated with Rapamycin (0.5 µM) or A/C Heterodimeriser (1 µM) over the course of 14 days. Medium with drugs was refreshed every 2 days. Confluence was analysed using an Incucyte time-lapse microscope. Representative of n = 3 biological replicates. FKBP12 - 12-kDa FK506-Binding Protein; FRB* - FKBP12-Rapamycin Binding Domain, T2098L; a.u. - arbitrary unit

Leveraging the spatiotemporal control of β-catenin and CSNK1D proximity enabled by the FKBP12^F36V^-FRB* dimeriser system, we next performed bulk RNAseq to gain systematic and unbiased insights into differential gene expression following treatment of cells with the A/C Heterodimeriser Rapalog for 8 h and 16 h (**Fig 4A**). Principal component analysis confirmed that variance between samples and biological replicates stemmed mainly from type and time of treatment (**Fig 4B**) and as expected, treatment with the A/C Heterodimeriser induced a robust downregulation of many known Wnt/β-catenin target genes at both time points, including *ASCL2*, *AXIN2*, *NKD1*, *RNF43, LEF1, ZNFR3* (**Fig 4C,D,E**). Accordingly, GO term analysis of downregulated differentially expressed genes included ‘Canonical Wnt signalling pathway’ as a driver term at both time points (**Fig 4F**) and the Hallmark gene set ‘Wnt β-catenin signalling’ was significantly depleted after A/C Heterodimeriser treatment in gene set enrichment analysis (**Fig 4G**).

**Fig. 4:**
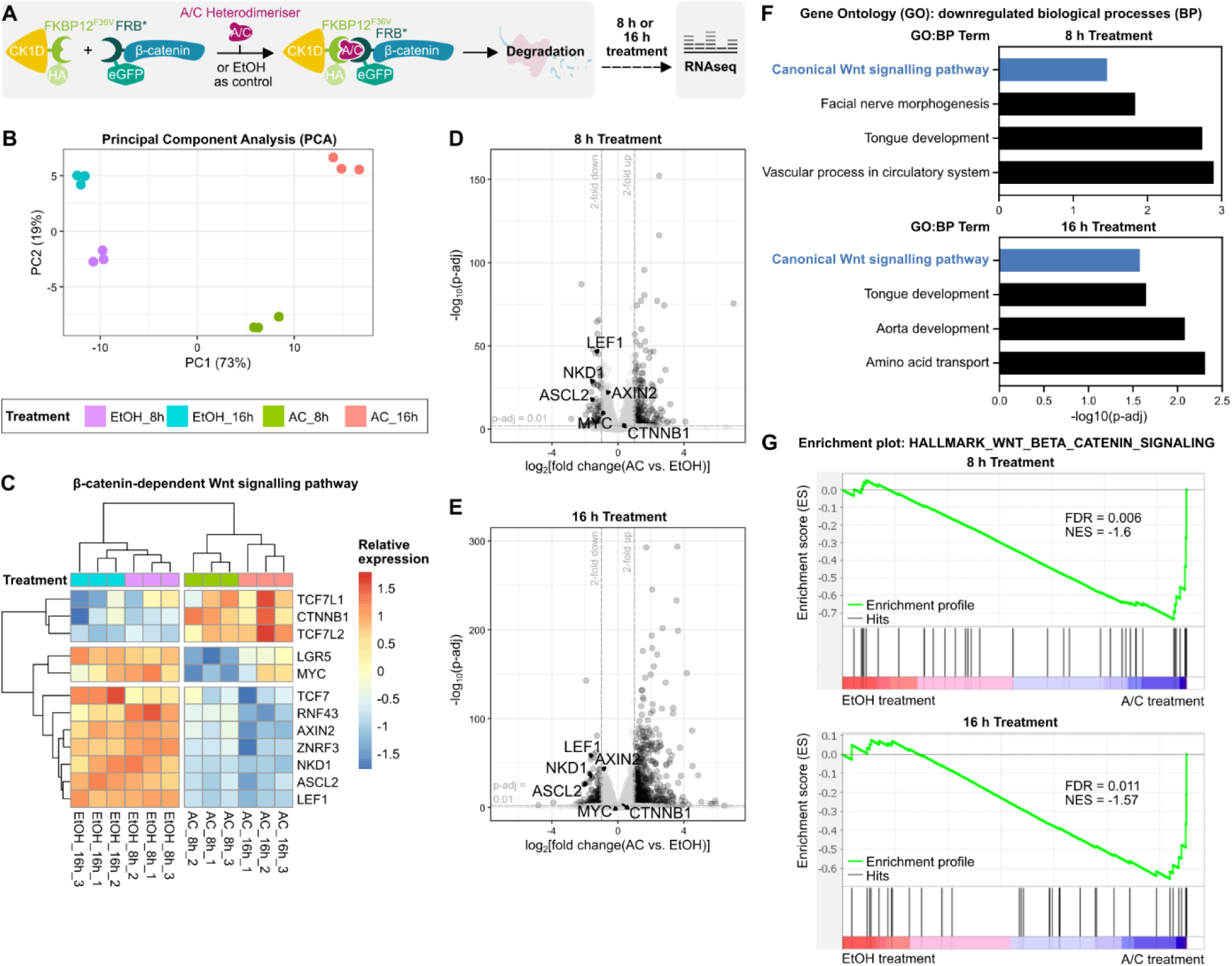
RNA-sequencing confirms specific regulation of Wnt signalling after CSNK1D-dependent degradation of β-catenin. **A)** Experimental outline of bulk RNA-sequencing (RNAseq) experiment: double tagged DLD-1 CTNNB1^eGFP-FRB*/eGFP-FRB*^ cells (clone #43) were infected with virus expressing CSNK1D-FKBP12^F36V^-HA and selected with puromycin for 2 days. Then, cells were treated with A/C Heterodimeriser (1 µM) or equivalent volumes of EtOH for 8 h or 16 h before RNA isolation for differential gene expression analysis. **B)** Principal component analysis (PCA) plot confirmed that variance between RNAseq samples of three biological replicates stemmed mainly from type and time of treatment. **C)** 8 h and 16 h treatment with the A/C Heterodimeriser induced a robust downregulation of Wnt/β-catenin target genes. Heatmap of scaled expression of genes associated with the β-catenin dependent Wnt signalling pathway in control (EtOH treated) or A/C Heterodimeriser treated samples. **D, E)** Volcano plots depicting differentially expressed genes at 8 h or 16 h after A/C vs. EtOH control treatment. Significantly regulated genes are shown in dark grey with Padj ≤ 0.01 and fold change > 2. **F)** Gene Ontology (GO) biological processes (BP) differentially downregulated in A/C Heterodimersier vs. EtOH control treated samples at 8 h and 16 h included ‘Canonical Wnt signalling pathway’ as a driver term. **G)** Enrichment plots depict depletion of genes contained in the human gene set ‘HALLMARK_WNT_BETA_CATENIN_SIGNALING’ (M5895) in A/C Heterodimeriser vs. EtOH control treated samples at 8 h and 16 h. ES - Enrichment score; NES – normalised enrichment score; FKBP12 - 12-kDa FK506-Binding Protein; FRB* - FKBP12-Rapamycin Binding Domain, T2098L

We conclude that forced proximity between β-catenin and CSNK1D induced by a small molecule leads to β-catenin degradation, downregulation of target genes and to inhibition of colorectal cancer cell proliferation.

### Targeted degradation of β-catenin by CSNK1D is kinase-activity and UPS-dependent

We were next interested to further understand the mechanism by which forced proximity of CSNK1D to β-catenin leads to its degradation. We first tested whether the kinase activity of CSNK1D was required. We showed that expression of two kinase-dead CSNK1D mutants, CSNK1D^K38R^ and CSNK1D^E247K/L252P^ (Liu et al., 2019; Xu et al., 2019), did not induce proximity dependent degradation of β-catenin (**Fig 5A**). We next probed the requirement of various cellular degradation pathways using chemical inhibitors. eGFP/eBFP2-β-catenin DLD-1 cells expressing NLuc-, CSNK1D- or SIAH2-vhhGFP were treated with MG132 (a proteasome inhibitor), TAK243 (an inhibitor of E1 ubiquitin activating enzyme), MLN-4924 (an inhibitor of neddylation) or Bafilomycin A (an inhibitor of lysosomal degradation) and cell lysates were probed using Western blotting to monitor β-catenin levels (**Fig 5B**). Inhibition of the UPS with either MG132, TAK243 or MLN-4924 partially rescued CSNK1D-induced degradation of β-catenin whereas Bafilomycin A had no appreciable effect. MG132 and TAK243 also rescued SIAH2-dependent degradation of β-catenin, but MLN-4924 did not, as SIAH2 does not require neddylation of a Cullin scaffolding protein for activity. Importantly probing for phospho-β-catenin levels, with an antibody recognizing phospho-threonine T41 and phospho-serine S45, showed its accumulation after inhibition of the UPS (**Fig 5B**). Together these results are consistent with β-catenin first getting phosphorylated following induced proximity with CSNK1D as a step required for degradation by the UPS.

**Fig. 5:**
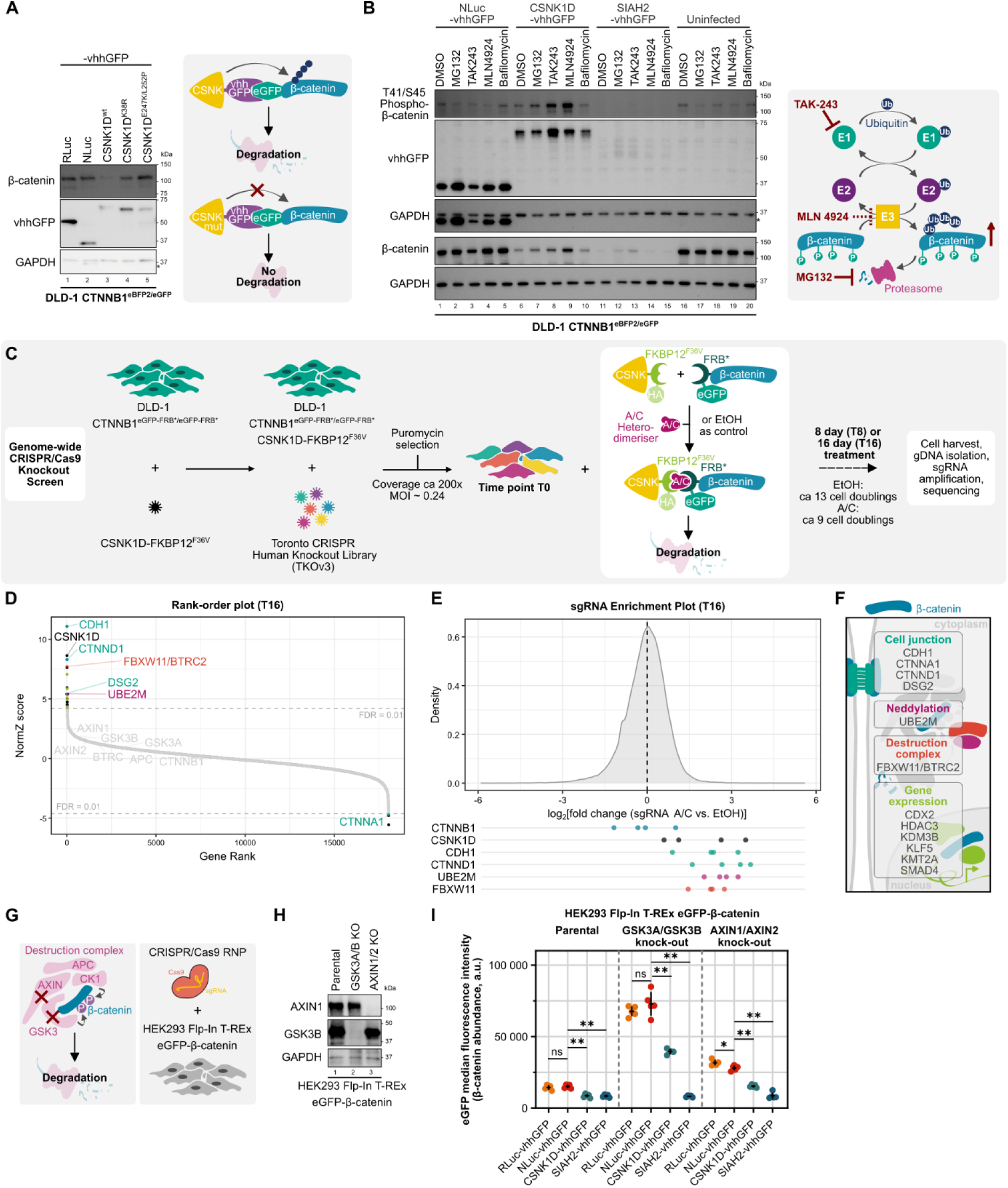
CSNK1D degrades β-catenin through a kinase- and Ubiquitin-Proteasome System-dependent mechanism. **A, B)** CSNK1D-dependent degradation of β-catenin is kinase activity and Ubiquitin-Proteasome System (UPS) dependent. DLD-1 CTNNB1^eBFP2/eGFP^ cells (clone #28) were infected with virus expressing the indicated -vhhGFP fusion proteins, selected with puromycin for 2 days and then analysed or further treated. **A)** Kinase-activity dead mutants of CSNK1D do not degrade eBFP2/eGFP-β-catenin. Representative Western Blot of n = 3 biological replicates. **B)** Inhibitors of the UPS rescue CSNK1D-dependent degradation of β-catenin and lead to accumulation of T41/S45 phosphorylated β-catenin. After selection, cells were treated with DMSO (0.1 %), the proteasome inhibitor MG132 (10 µM), the E1 inhibitor TAK-243 (1 µM), the inhibitor of neddylation MLN-4924 (10 µM) or Bafilomycin (1 µM) for 18 h before harvest and analysis by Western Blot. **C, D, E, F)** Genome-wide CRISPR/Cas9 knock-out fitness screen identifies regulators of CSNK1D-dependent β-catenin degradation. Highlighted are genes involved in cell-cell contacts (green), transcriptional regulation (light green), neddylation (pink), the SCF^β-TrCP^ E3 ubiquitin ligase complex (red, FBXW11 is also known as BTRC2), and others (black). **C)** Schematic of positive selection screen in double tagged DLD-1 CTNNB1^eGFP-FRB*/eGFP-FRB*^ cells (clone #43) with stable expression of CSNK1D-FKBP12^F36V^-HA. After puromycin selection, cells were split into three replicates per condition and treated with A/C Heterodimeriser (1 µM) or equivalent volumes of EtOH. Media and compounds were refreshed every 2 days. **D)** Rank order plot depicting the gene level summary of sgRNAs that regulate CSNK1D-dependent β-catenin degradation identified at Endpoint (T16). Black dots represent regulators with FDR < 0.01. **E)** overall sgRNA distribution and individual sgRNA enrichment at T16 of the highlighted genes, summarised in **F)**. **G, H, I)** GSK3 and AXIN are dispensable for CSNK1D-induced degradation of β-catenin. **G)** HEK293 Flp-In T-REx GFP-CTNNB1 cells were transfected with CRISPR ribonucleoproteins (RNPs) to generate GSK3A/GSK3B or AXIN1/AXIN2 double knock-out (KO) cell lines. **H)** Western blot confirms gene knock-out. **I)** HEK293 Flp-In T-REx cells stably expressing eGFP-β-catenin under a TET-inducible promoter were transfected with the indicated - vhhGFP fusion proteins and eGFP expression was measured by flow cytometry as read-out for β-catenin abundance. Individual data points of four or five independent experiments are shown with mean and standard deviation. A minimum of 4,500 cells per condition were recorded in the final gate. Statistical significance was assessed by one-way ANOVA followed by Dunnett’s multiple comparisons test comparing all -vhhGFP groups to NLuc-vhhGFP. * Padj ≤ 0.05, ** Padj ≤ 0.0001, ns = not significant. For Western blots: * signal from previous staining; kDa = Kilodalton; GAPDH served as loading control; RLuc - Renilla luciferase; NLuc - NanoLuc luciferase; vhhGFP - nanobody binding to eGFP and eBFP2; a.u. - arbitrary unit

To systematically determine which proteins are involved in CSNK1D-dependent degradation of β-catenin, we performed a genome-wide CRISPR/Cas9 suppressor screen in eGFP-FRB*-β-catenin DLD1 cells also expressing CSNK1D-FKBP12^F36V^ in the presence of the Rapalog A/C Heterodimeriser (**Fig 5C**). Since treatment of these cells with the Rapalog led to inhibition of cell proliferation (**Fig 3D**), knock-out of genes that rescue β-catenin protein abundance are predicted to also rescue cell proliferation and to be positively selected. Predictably, CSNK1D itself scored as one of the top hits. Further validating our approach, KMT2A and ZCCHC14 are hits that were also identified in previously published genome-wide CRISPR/Cas9 screens for Wnt/β-catenin signalling in colon cancer cells (Wan et al., 2021). Other significant hits after 8 days and 16 days of treatment drew a comprehensive picture of β-catenin in colorectal cancer cells (**Fig 5D,E,F, Suppl Fig 5A**). These included β-catenin interaction partners at the plasma membrane (e.g. CDH1, CTNND1, DSG2), potentially because interference with cell junction integrity released β-catenin from the membrane and increased cytoplasmic and nuclear β-catenin pools available for transcriptional regulation. Other hits control gene expression and possibly interact with β-catenin in the nucleus or regulate epigenetic mechanisms (e.g. CDX2, HDAC3, SMAD4). Importantly, top hits also included components of the neddylation machinery and the Skp1-Cullin-F-box-protein (SCF) (FBXW11/BTRC2) E3 ubiquitin ligase complex, confirming their requirement for CSNK1D-dependent β-catenin degradation through induced-proximity. FBXW11 is one of two human paralogous β-TrCP family members, which act as substrate adaptors in SCF E3 ubiquitin ligase complexes in the context of the β-catenin targeting destruction complex (Maurice and Angers, 2025; Wu et al., 2003).

We found curious that the β-catenin destruction complex components GSK3α/β or AXIN1/2 were not identified in our CRISPR screen, possibly because of the functional redundancy between the homologs. To directly test their requirement for CSNK1D-dependent β-catenin degradation, we created *GSK3A/GSK3B* and *AXIN1/AXIN2* double knock-out HEK293 Flp-In T-REx cell lines that express eGFP-β-catenin under a tetracycline inducible promoter and measured eGFP expression using flow cytometry after transfection with CSNK1D-, SIAH2, RLuc- or NLuc-vhhGFP (**Fig 5G,H, Suppl Fig 5B**). eGFP-β-catenin levels were elevated in both double knock-out cell lines, as predictably explained by disabling destruction complex function (**Fig 5I**). Both CSNK1D- and SIAH2-vhhGFP efficiently degraded eGFP-β-catenin in the absence of GSK3a/b or AXIN1/2 (**Fig 5I**). Similar results were obtained in eGFP/eBFP2-β-catenin DLD-1 cells where siRNA-mediated knockdown of GSK3a/b did not interfere with CSNK1D-vhhGFP mediated degradation while knockdown of *BTRC* and *FBXW11* (β-TrCP1/2) impeded CSNK1D-induced degradation of β-catenin and led to accumulation of its phosphorylated form (**Suppl Fig 5C**). We conclude that induced proximity to CSNK1D and recruitment of a functional SCF complex containing β-TrCP2 are sufficient to respectively induce β-catenin phosphorylation and degradation thereby bypassing the need for other destruction complex components.

### Induced proximity to CSNK1D degrades some but not all oncogenic β-catenin mutants

In the context of the destruction complex, β-catenin is first phosphorylated by CSNK1A1 at S45 and then by GSK3a/b at T41, S37, and S33 (Amit et al., 2002; Liu et al., 2002). Phosphorylated S37 and S33 are the residues underlying binding to β-TrCP1/2, a substrate specific adapter for the SCF ubiquitin ligase complex, which ubiquitinates β-catenin leading to its proteasomal degradation (Aberle et al., 1997; Orford et al., 1997; Wu et al., 2003) (Fig 6A). In addition to the very frequent *APC* mutations, mutations within the β-catenin S45, T41, S37, S33 phospho-degron are also found in different cancers as they also impair β-catenin degradation leading to ligand-independent activation of the pathway and to uncontrolled cell proliferation (Kim and Jeong, 2019; Krishna et al., 2026). As we showed that promoting the proximity of CSNK1D to β-catenin is sufficient to promote its phosphorylation and degradation and bypassed other destruction complex components, we were next interested to map the exact β-catenin phospho-degron residues that are required for degradation in this context. These results are also important to understand which ones of the various oncogenic β-catenin mutations would still be subject to degradation, following induced proximity of CSNK1D, to guide the use of future PROTAC molecules leveraging this concept. To do this, we generated a series of inducible isogenic cell lines that express eGFP-β-catenin variants under a tetracycline inducible promoter and measured eGFP expression using flow cytometry following expression of CSNK1D-, SIAH2, RLuc- or NLuc-vhhGFP (**Fig 6B**). Wild-type β-catenin was potently degraded by CSNK1D- and SIAH2-vhhGFP, as expected. We then tested β-catenin phospho-mutants where S45, T41, S37, or S33 (or various combinations thereof) were deleted or replaced by alanine (A) and various combinations thereof. Notably, the eGFP-β-catenin mutants were all expressed at higher levels than wild-type β-catenin reflecting stabilization resulting from impaired regulation by the destruction complex machinery (**Fig 6B**). Expression of SIAH2-vhhGFP robustly degraded all eGFP-β-catenin variants in this assay. Strikingly, induced proximity to CSNK1D-vhhGFP degraded both the S45del and T41A eGFP-β-catenin mutants, but no other tested mutants (**Fig 6B**). We conclude that phosphorylation of S45 or T41 is dispensable in the context of CSNK1D-dependent degradation and that oncogenic β-catenin mutants with alterations at these positions, which are the most common *CTNNB1* mutations found in cancer patients (Krishna et al., 2026), would still be susceptible for degradation.

**Fig. 6:**
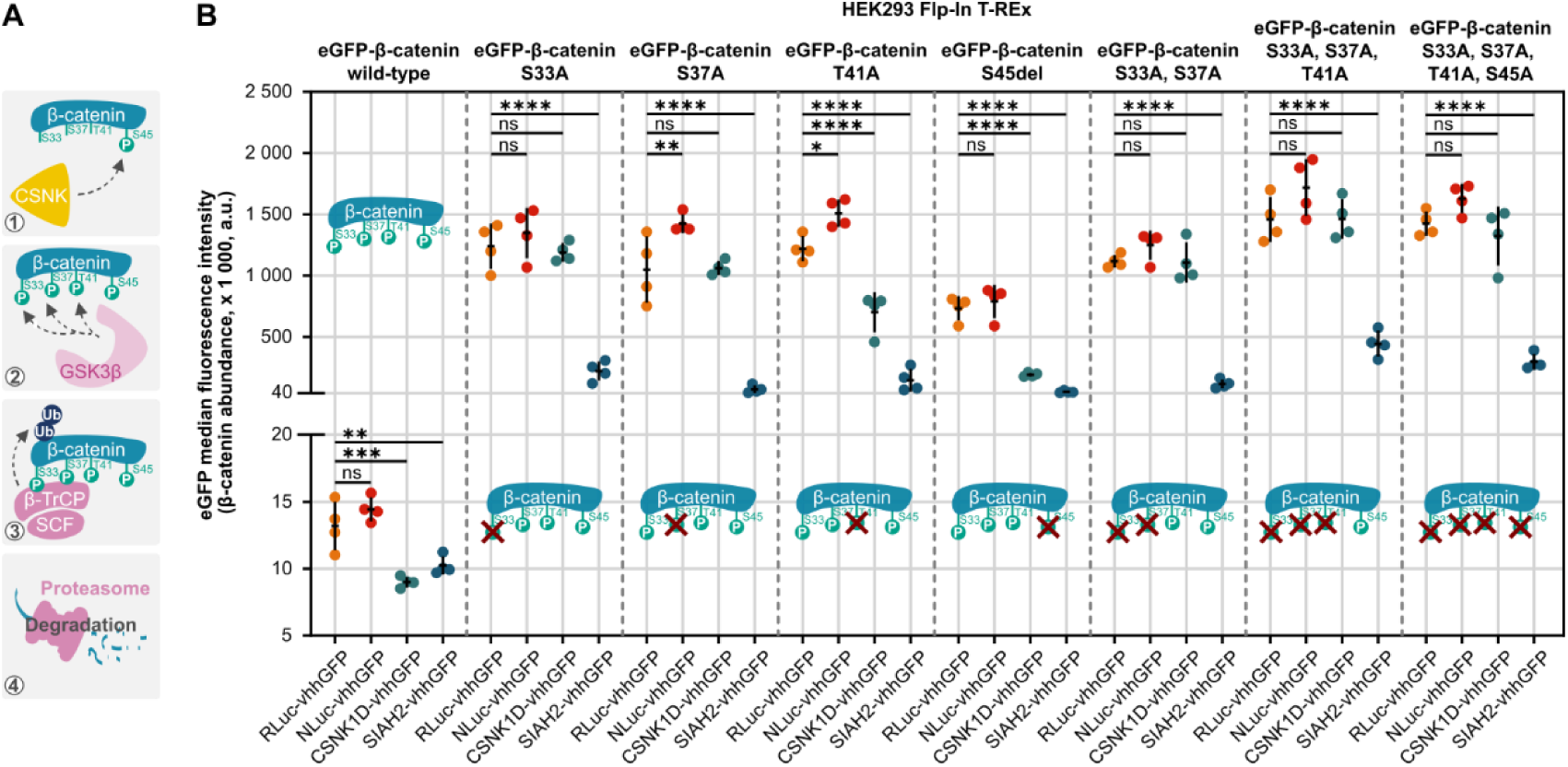
Induced proximity to CSNK1D degrades oncogenic β-catenin mutants. **A, B)** In the context of the destruction complex, β-catenin is first phosphorylated at serine S45 by CSNK1 and then by GSK3 on threonine T41, S37, and S33. Phosphorylated S33 and S37 and surrounding amino acids are bound by β-TrCP, leading to β-catenin ubiquitinylation and degradation by the proteasome. Mutations of any of these positions have been found in the context of cancer and protect β-catenin from degradation by the endogenous destruction complex. However, phospho-dead β-catenin mutants are degraded efficiently by induced proximity to CSNK1D and SIAH2 as long as the β-TrCP binding sites are not mutated. HEK293 Flp-In T-REx cells stably expressing eGFP-β-catenin wild-type (wt) or phospho-dead mutants under a TET-inducible promoter were transfected with the indicated -vhhGFP fusion proteins and eGFP expression was measured using flow cytometry as read-out for β-catenin abundance. Individual data points of four independent experiments are shown with mean and standard deviation. A minimum of 5,500 cells per condition were recorded in the final gate. Statistical significance was assessed by one-way ANOVA followed by Dunnett’s multiple comparisons test comparing all - vhhGFP groups to RLuc-vhhGFP. * Padj ≤ 0.05, ** Padj ≤ 0.01, *** Padj ≤ 0.001, **** Padj ≤ 0.0001, ns = not significant. RLuc - Renilla luciferase; NLuc - NanoLuc luciferase; vhhGFP - nanobody binding to eGFP and eBFP2; a.u. - arbitrary unit

### Promoting degron formation through kinase recruitment extends the PROTAC toolbox

Our results suggest that restoring phospho-degron formation through forced recruitment of a kinase to a substrate can restore its degradation in a context where the process is malfunctioning, such as in cancer. To assess the generality of this concept we turned to the transcriptional regulator SNAI1, which is involved in epithelial-to-mesenchymal-transition and undergoes a similar phospho-dependent degradation as β-catenin. Following a priming phosphorylation event by CSNK1, SNAI1 is further phosphorylated by GSK3α/β leading to degradation by β-TrCP-mediated ubiquitination (Xu et al., 2010). We expressed eGFP-SNAI1 together with various vhhGFP-tagged fusion proteins and monitored eGFP-SNAI1 expression by flow cytometry. Strikingly, eGFP-SNAI1 was degraded in the presence of CSNK1D-vhhGFP to the same extent as with CRBN-vhhGFP used here as a positive control (CRBN is the most widely used E3 ligase component used in PROTACs (Hinterndorfer et al., 2025; King et al., 2025)). The kinase dead mutant CSNK1D-vhhGFP led to a small stabilization of eGFP-SNAI1 pointing at a kinase-dependent degradation mechanism in this context as well (**Fig 7A**). In summary, we propose a novel application for TPD, where forced recruitment of a kinase to a substrate induces neo phospho-degron formation on a target protein of interest, leading to its recognition by endogenous E3 ubiquitin ligases, catalysing ubiquitination and degradation by the proteasome. Although degradation of a wide variety of proteins may be possible using this approach, we anticipate that it will be particularly useful to target substrates of SCF complexes in contexts where their degradation is defective such as in cancer (**Fig 7B**).

**Fig. 7:**
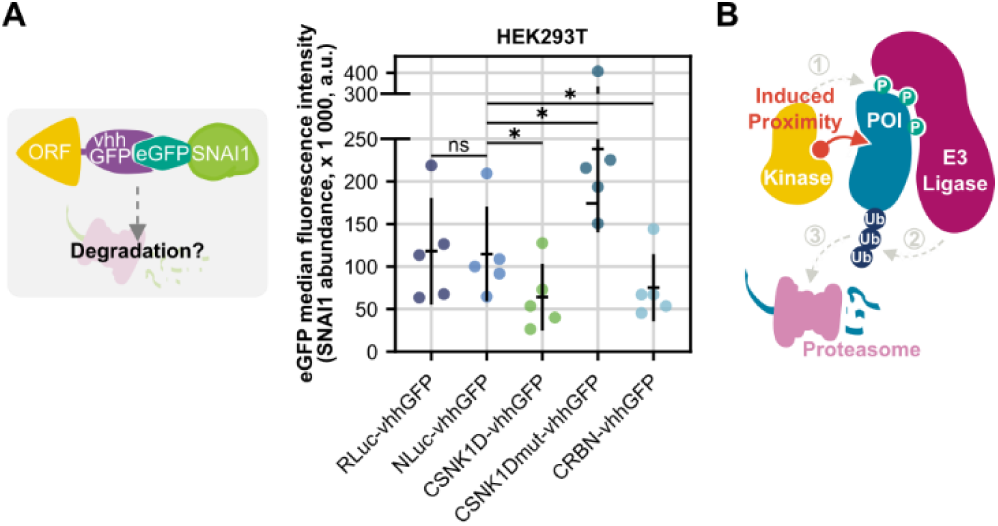
eGFP-SNAI1 is degraded by induced proximity to CSNK1D. **A)** eGFP-SNAI1 is degraded by CSNK1D-vhhGFP in a kinase dependent mechanism. HEK293T cells were transfected with eGFP-SNAI1 and -vhhGFP tagged fusion proteins. 24 h later, GFP expression was measured using flow cytometry as read-out for SNAI1 abundance. Individual data points of five independent experiments are shown with mean and standard deviation. A minimum of 2,500 cells per condition were recorded in the final gate. Statistical significance was assessed by Repeated Measures one-way ANOVA followed by Dunnett’s multiple comparisons test comparing all -vhhGFP groups to NLuc-vhhGFP. * Padj ≤ 0.05, ns = not significant. CSNK1Dmut – CSNK1D^E247K/L252P^; RLuc - Renilla luciferase; NLuc - NanoLuc luciferase; vhhGFP - nanobody binding to eGFP and eBFP2; a.u. - arbitrary unit **B)** Schematic depiction of kinase-dependent targeted protein degradation: Induced proximity to kinases leads to phosphorylation of the protein of interest (POI). This phospho-degron is recognised by an E3 ubiquitin ligase which labels its substrate with Ubiquitin (Ub), resulting in proteasomal target protein degradation.

## DISCUSSION & OUTLOOK

Here, we used orthogonal proteome-wide screening strategies to identify induced-proximity regulators of β-catenin protein abundance and colon cancer cell proliferation (Since β-catenin is essential for their growth). We revealed potent proximity-dependent regulators of β-catenin, many of which were E3 ubiquitin ligases, confirming the efficiency of our screening strategy. Some of these effectors were previously described to degrade other substrates (Poirson et al., 2024) and are most likely potent, but not necessarily β-catenin specific degraders. More interestingly, we found candidates that had not been explored in the context of TPD, namely members of the kinase family CSNK1 and the RING E3 ubiquitin ligase SIAH2. The induced turnover of β-catenin upon recruitment of CSNK1D resulted in sustained and phenotypically relevant downregulation of β-catenin protein expression that inhibited colorectal cancer cell growth. Further functional and mechanistic characterisation confirmed that β-catenin degradation by CSNK1D was proximity-, kinase activity-, and proteasome-dependent and required activity of the E3 ubiquitin ligase adapter FBXW11/BTRC2. Thus, we proposed the formation of a neo phospho-degron on β-catenin as underlying degradation mechanism.

While our proof-of-concept study relied on antibody- and small-molecule-mediated proximity using various protein tags, to fully leverage this concept it will now be important to design cell permeable bifunctional molecules that will promote the recruitment of CSNK1D with β-catenin. The availability of direct small molecule β-catenin binders may currently be a limitation to such approach (McCoy et al., 2022). Similarly, non-inhibitory CSNK1D binders do not exist. However, recent developments in the field suggest it may be feasible to employ kinase inhibitors for a ‘catch and release-mechanism’ (Sarott et al., 2024), or cysteine-based group transfer chemistry (Pergu et al., 2023) to turn existing orthosteric kinase inhibitors into potent binders for induced-proximity purposes.

Many compounds have been developed that interfere with either components of the β-catenin destruction complex or the transcriptional machinery associated with β-catenin, but to date, no drugs have been approved to treat diseases characterized with hyperactive Wnt signalling (Maurice and Angers, 2025; Parsons et al., 2021; Wang et al., 2021). A VHL-based PROTAC molecule was developed that binds β-catenin via a small, Axin-derived, stapled peptide but only showed moderate activity in the high micromolar range in validation experiments (Grossmann et al., 2012; Liao et al., 2020). Interestingly, our screen results indicate that VHL does not seem to be a very potent degrader of β-catenin highlighting the benefit to identify induced-proximity candidates in unbiased methods. In another approach, high-throughput screening and rational design led to the identification of a molecular glue that directly stabilizes the interaction between a mutant β-catenin variant and its natural E3 ligase target receptor, β-TrCP (Simonetta et al., 2019). While this discovery laid important groundwork for the prospective design of small molecule degraders, this molecular glue fails to interact with the most frequent β-catenin mutants found in cancer. Importantly, we showed that CSNK1D-mediated degradation is more versatile and efficiently targets the frequent S45 and T41 cancer causing mutations. A monovalent covalent binder of β-catenin that destabilizes and degrades β-catenin was also recently identified (Gowans et al., 2024). While the precise mechanism of action remains to be explored, improved potency and stability might be needed to sustain durable β-catenin down regulation and persistent biological effects.

Kinases and protein phosphorylation have previously been leveraged in other induced proximity contexts to modify cellular effects. For instance, signalling pathways were modulated through synthetic tyrosine phosphorylation (Pergu et al., 2023; Shoba et al., 2022; Siriwardena et al., 2020) and apoptosis was triggered by repurposing transcriptional kinases (Sarott et al., 2024). Other recent studies used chemical-induced dephosphorylation to regulate kinase signalling (Simpson et al., 2023; Sun et al., 2024). However, to our knowledge, the current study represents the first example where forced recruitment of a kinase to a protein promotes its degradation. We envision that this concept will be extended beyond β-catenin to modulate the activity of other cellular proteins that depend on phosphorylation as a recognition signal for their degradation. One particularly attractive class of substrates are those regulated by phospho-degrons recognized by F-box proteins as part of SCF ubiquitin ligase complexes. To showcase this potential, we demonstrated that we could promote the degradation of the Zinc finger protein SNAI1 by forcing its recruitment to CSNK1D (**Fig 7**). Therefore, we propose that promoting phosphorylation of proteins will expand the chemical toolkit for the design of targeted proximity degradation modalities.

Another strong degrader of β-catenin we characterised here was the RING E3 ubiquitin ligase SIAH2. SIAH2 had been associated with β-catenin turnover before, mainly in the context of Wnt5a-induced inhibition of Wnt/β-catenin signalling and hypoxia in melanoma (O’Connell et al., 2013; Topol et al., 2003). However, it remained unclear if SIAH2 was directly targeting β-catenin for ubiquitination or whether this effect was mediated through alternative substrates. Importantly, direct ubiquitination of non-phosphorylated β-catenin was shown for SIAH1, the closely related homolog of SIAH2, in biochemical assays and as part of the p53-inducible response in cells (Dimitrova et al., 2010; Matsuzawa and Reed, 2001). The high sequence homology between SIAH1 and SIAH2, and our results presented here, indicate that SIAH2 indeed directly and potently regulates β-catenin independent of its phosphorylation status and has potential for TPD applications.

While CSNK1A1 (Casein kinase I α) is generally considered to be the endogenous priming kinase of β-catenin within the destruction complex in healthy cells (Liu et al., 2002; van Kappel and Maurice, 2017), it was shown that other members of the CSNK1 family can phosphorylate β-catenin, at least in biochemical studies (Amit et al., 2002). It is curious that CSNK1A1 was not identified in our induced proximity screens, while we recovered other members of the family (CSNK1D, CSNK1E and CSNK1G2) in at least one of the screens. A possible explanation may be the smaller size of CSNK1A1 that consists mainly of a kinase domain when compared to other family members that also have a non-catalytic C-terminal domain [human CSNK1A1 (UniProt ID P48729): 337 AA, human CSNK1D (UniProt ID P48730): 415 AA]. As all effectors were tagged at the C-terminus with vhhGFP in our screens, it is conceivable that the smaller CSNK1A1 cannot reach its designated phosphorylation sites on β-catenin within the effector/target complex. Our results also clearly suggest that phosphorylation of β-catenin at S33 and S37 is required for SCF^β-TRCP^ mediated degradation (**Fig 6A**) and that forcing the recruitment of CSNK1D promotes the degradation of all oncogenic β-catenin mutants as long as these two sites are not mutated. These results are consistent with our suppressor screen results that revealed the requirement of the E3 ubiquitin ligase adapter protein FBXW11 (also known as β-TrCP2) for CSNK1D-mediated degradation. This approach should therefore be efficient in the vast majority of Wnt-HIGH cancers in which β-catenin is stabilised as a consequence of *APC* mutations or the most frequent mutations affecting the β-catenin phospho-degron (missense mutations of residues S45 or T41 of β-catenin account for 25% and 27% of all mutations in this hotspot region, respectively (Krishna et al., 2026). A bifunctional molecule mediating CSNK1D and β-catenin proximity would therefore act to chemically bypass the activity of the destruction complex to mediate β-catenin degradation and re-establish homeostatic transcriptional activity levels. Collectively our data offers a novel strategy to target pathogenic high Wnt-β-catenin signalling in diseases such as cancer when the pathway is activated as a result of disarming mutations within the destruction complex or stabilizing mutations within β-catenin itself.

## MATERIAL & METHODS

### Cell Culture

Colorectal adenocarcinoma cell line DLD-1 (American Type Culture Collection, ATCC, CCL-221) and derived cell lines were cultured in RPMI-1640 medium (Gibco, Thermo Fisher Scientific, 11875119) supplemented with 10% Fetal Bovine Serum (FBS, Gibco, Thermo Fisher Scientific, 12483020) and 1% Penicillin-Streptomycin (Gibco, Thermo Fisher Scientific, 15140122). HEK293T (ATCC, CRL-3216) and HEK293 Flp-In TRex (Invitrogen, Thermo Fisher Scientific, R78007) cells were cultured in DMEM with 4.5 g/L Glucose, L-Glutamine & Sodium Pyruvate (Wisent Bioproducts, 319005 CL) supplemented with 10% FBS and 1% Penicillin-Streptomycin, as above. All cell lines were regularly confirmed to be mycoplasma negative using the MycoAlert® Mycoplasma Detection Kit (Lonza, LT07-118) and authenticated using Short Tandem Repeat (STR) profiling.

### Generation of endogenously tagged DLD-1 cell lines

To endogenously tag β-catenin at its N-terminus with eBFP2/eGFP, eGFP-FRB*, or HaloTag, 360 000 DLD-1 cells were electroporated using the Neon Transfection System 10 µl Kit (Invitrogen, Thermo Fisher Scientific, MPK1025) following the manufacturer’s instructions with 1.5 µg CleanCap Cas9 mRNA (TriLink Biotechnologies, L-7206), 1.2 µg Universal sgRNA (5’-GGGAGGCGUUCGGGCCACAG-3’, std mod SpCas9 template, Synthego), 1.2 µg CTNNB1 sgRNA (5’-UGAGUAGCCAUUGUCCACGC-3’, std mod SpCas9 template, Synthego) and 1.2 µg donor plasmid (Settings: 1530 V, 20 ms, 1x pulse). Donor plasmids were cloned so that UgRNA cut sites framed the tag to be inserted into the genome and 48 bp homology arms (Welker et al., 2021; Wierson et al., 2020). See supplemental data for sequences of constructs for insertion. The eBFP2/eGFP-tags were inserted into the DLD-1 βcatenin-DEADPOOL cell line (Mis et al., 2020), while GFP-FRB* and HaloTag were inserted into DLD-1 Flp-In cells.

Cells were left to recover for 5 days and then analysed for fluorophore expression/tag integration using Fluorescence Activated Cell Sorting (FACS). Fluorescent cells were single cell sorted into 96 well plates [Flow buffer: 1 mM EDTA (BioShop Canada, EDT001), 25 mM HEPES (Gibco, Thermo Scientific, 15630080), 1% BSA in PBS]. To detect HaloTag insertion, cells were labelled with HaloTag NanoBRET 618 Ligand (Promega, G980A) before sorting. Clonal cell lines were expanded and characterised by Western blotting to detect a size shift of β-catenin protein and sequencing of genomic DNA to confirm seamless integration of the tags. To do so, genomic DNA from ca. 1×10^6 cells was isolated using the PureLink Genomic DNA Mini Kit (Invitrogen, Thermo Fisher Scientific, K182002) and the transcriptional start site of CTNNB1 was amplified using primers spanning the tag integration site (fwd, 5’-GTGGTTGAGGTGTCTGGAGG-3’, rev: 5’-ATTCCTGCTGGTGGCTTGTT-3’) and the GoTaq Green Master Mix (Promega, M7122) according to the manufacturer’s instructions. In brief, 500 ng to 1 µg of genomic DNA was used as template in a 25 µl reaction and the following cycle protocol: 95 °C – 3 min (initial denaturation), 95 °C – 30 sec (denaturing), 58 °C – 30 sec (annealing), 72 °C – 1 min 20 (extension, ca 1 min per 1 000 bp), 72 °C – 5 min (final extension), with a total of 30 cycles. Amplicons were separated on a 1.5% agarose gel, gel extracted using the E.Z.N.A. Gel Extraction Kit (Omega Bio-tek, D2500-02) and analysed using Sanger sequencing.

For eGFP-FRB*- and HaloTag-CTNNB1 DLD-1, several clones in which both alleles had been tagged were identified after a single round of electroporation (double tagged cell lines). To create the eBFP2/eGFP-tagged cell line, the first round of electroporation with the eGFP donor plasmid resulted in the heterozygous integration of eGFP and a 12 bp deletion at the transcriptional start site of the untagged allele. However, this allele was still expressed, and a second round of electroporation with an adjusted sgRNA and the eBFP2 donor plasmid repaired the deletion and inserted eBFP2 in parallel to eGFP, resulting in the single cell clones #28, #84, and #92.

### Stable HEK293 Flp-In T-REx GFP-CTNNB1 cell lines and protein degradation analysis

To generate isogenic cell lines with stable, tetracycline-inducible expression of eGFP-β-catenin variants, HEK293 Flp-In T-REx (Flp-In T-REx 293 Cell Line, Invitrogen, Thermo Fisher Scientific, R78007) were transfected with plasmid pOG44 and respective pcDNA5/FRT/TO construct (see Supplement) at a DNA ratio of 3:1 using Lipofectamine 2000 Transfection Reagent (Invitrogen, Thermo Fisher Scientific, 11668027) according to the manufacturer’s recommendations (6 µl Lipofectamine 2000 per 1 µg of DNA). Two days after transfection, selection with 50 µg/ml Hygromycin B (BioShop Canada Inc., HYG002) was started and continued for approximately 2 weeks until all control cells without Hygromycin resistance had died.

HEK293 Flp-In T-Rex eGFP-CTNNB1 GSK3A/GSK3B or AXIN1/AXIN2 double knock-out cell lines were generated by seeding 80 000 cells of the parental cell line per well of a 24 well plate in 500 µl growth medium. The next day, 2 µl of two human Gene Knockout Kits (3 pmol/µl each, EditCo, GKO-HS1-000-0-1.5n-0-0) were complexed for 5 min at room temperature with 1 µl TrueCut Cas9 Protein v2 (Invitrogen, Thermo Fisher Scientific, A36496) in 25 µl Opti-MEM (Gibco, Thermo Fisher Scientific, 11058021). At the same time, 1 µl Lipofectamine 3000 Transfection Reagent (Invitrogen, Thermo Fisher Scientific, L3000008) were diluted in 25 µl Opti-MEM, mixed briefly, incubated for 1 min at room temperature and then combined with the complexed Cas9 ribonucleoproteins (RNPs). After another 10 min incubation at room temperature, the mixture was added dropwise to the cell. Cells were left to recover and expand for 5 days. Then, eGFP-CTNNB1 expression was induced with tetracyclin (Bioshop Canada, TET701) and the top 5 % of strongest GFP expressing cells were bulk sorted at the University of Toronto Centre for Immune Analytics Core Facility (RRID: SCR_027612), as successful knock-out of GSK3A/GSK3B or AXIN1/AXIN2 should stabilize eGFP-β-catenin expression. The targeted DNA sequences were amplified by PCR and InDel percentage and knockout score were determined using the ICE (Inference of CRISPR Edits) tool (https://ice.editco.bio).

For experimental analysis, 1×10^6 HEK293 cells were seeded in 2 ml growth medium per well in 6 well plates and eGFP-β-catenin construct expression was induced with 1 µg/ml tetracycline (Bioshop Canada, TET701). One well per cell line was left untreated as control. The next day, cells were transfected with pcDNA3.1-[ORF]-GSlinker-vhhGFP-SV40-TagRFP or pcDNA3.1-[ORF]-GSlinker-NbALFA-GSlinker-3xMyc-tag-SV40-TagRFP expressing plasmids using Lipofectamine 2000 Transfection Reagent according to the manufacturer’s recommendations. In brief, per well, 1 µg plasmid DNA was diluted in 100 µl Opti-MEM (Gibco, Thermo Fisher Scientific, 11058021) and 3 µl Lipofectamine 2000 were diluted in 100 µl Opti-MEM. After 5 min incubation at room temperature, both solutions were combined, mixed gently and incubated for another 20 min at room temperature before being added drop-wise to the cells. 24 h later, eGFP expression as proxy for β-catenin abundance was analysed using flow cytometry. TagRFP expression was used to determine transfected cells and a TagRFP-GSlinker-3xFLAG-V5-tag only plasmid was used as control for gating.

### Plasmids

Gateway technology was used to clone entry clones picked from the hORFeome collection into pLX301-[ORF]-vhhGFP, pLX301-[ORF]-NbALFA-GSlinker-3xMyc-tag, or pLX301-[ORF]-FKBP12^F36V^-HA-tag destination vectors for lentivirus production and infection of DLD-1 cells.

For expression in HEK293 or HEK293T cell lines, entry clones were integrated in pcDNA3.1-[ORF]-GSlinker-vhhGFP-SV40-TagRFP or pcDNA3.1-[ORF]-GSlinker-NbALFA-GSlinker-3xMyc-tag-SV40-TagRFP.

Donor plasmids for endogenous tagging in DLD-1 and pcDNA5/FRT/TO constructs for stable integration into HEK293 Flp-In T-Rex cells, as well as plasmids for competition assays were cloned using standard methods and the sequences of inserts can be found in the Supplement.

The two kinase-dead CSNK1D variants CSNK1D K38R and E247K/L252P (Liu et al., 2019) were derived from the parental plasmid using the QuikChange II XL Site-Directed Mutagenesis Kit (Agilent Technologies, 200521) and the following primers: K38R (fwd: 5’-ggagaagaggttgccatcaggcttgaatgtgtcaaa-3’, rev: 5’-tttgacacattcaagcctgatggcaacctcttctcc-3’), E247K/L252P (fwd: 5’-aaaggctacccttccaaatttgccacatacccgaatttctgccgttc-3‘, rev: 5‘-gaacggcagaaattcgggtatgtggcaaatttggaagggtagccttt-3‘).

GFP-SNAI1 was cloned from a SNAI1 clone picked from the hORFeome collection (canonical uniport isoform) to make the pcDNA3.1-EGFP-SNAI1 plasmid.

### siRNA transfection

One day before transfection, DLD-1 cells were seeded in a 6 well plate in 2 ml growth medium (250 000 cells/well). The next day, 125 µl Opti-MEM (Gibco, Thermo Fisher Scientific, 31985070) were mixed with 6 µl Lipofectamine RNAiMAX Transfection Reagent (Invitrogen, Thermo Fisher Scientific, 13778075) and in parallel, 125 µl Opti-MEM were mixed with 3 µl (siGFP, siCTNNB1) or 6 µl (siBTRC1/2, siGSK3A, siGSK3B) siRNA (all siRNAs were diluted to 5 µM stock concentration). After incubation at room temperature for 2 min, the two volumes were combined, mixed again carefully and incubated at room temperature for another 5 min, then added dropwise to the cells (final siRNA conc 6.7 nM or 13.4 nM). Untransfected cells and cells incubated with transfection reagent only (mock transfection) were included as controls. Cells were harvested for analysis 48 h to 72 h after transfection.

### Lentivirus production and infection

Lentivirus was produced by transfecting 1×10^6 HEK293T cells in 2 ml growth medium per well of a 6 well plate with respective 1.25 µg pLX301 construct, 900 ng psPAX2 (Addgene, 12260) and 300 ng pVSV-G (Addgene, 8454) using Lipofectamine 2000 Transfection Reagent (Invitrogen, Thermo Fisher Scientific, 11668027) according to the manufacturer’s protocol. In brief, DNA was diluted in 100 µl Opti-MEM (Gibco, Thermo Fisher Scientific, 11058021) and 6 µl Lipofectamine 2000 were diluted in 100 µl Opti-MEM. After 5 min incubation at room temperature, both solutions were combined, mixed gently and incubated for another 25 min at room temperature before being added drop-wise to the cells. 6 h to 16 h after transfection, the medium was changed to 1.5 ml growth medium containing 1.1 g/100 ml Bovine Albumin Fraction V (Gibco, Thermo Fisher Scientific, 15260037). Two days after transfection, supernatant was filtered (0.45 μm filter) and used to infect cells or stored at - 80 °C.

In infect DLD-1, 600 000 cells per well of a 6-well plate were infected using 300 – 500 µl virus in 2 ml growth medium in the presence of 8 µg/ml Polybrene (Hexadimethrine bromide, Sigma, H9268). The next day, selection with 1.5 µg/ml puromycin (Gibco, Thermo Fisher Scientific, A1113803) was started and continued for 2 days, after which cells were analysed. To increase the selection pressure, cells were lifted after 1 day and re-seeded into the same well.

### RNA isolation, complementary DNA (cDNA) synthesis and reverse-transcription quantitative PCR (RT-qPCR)

For gene expression analysis, RNA was isolated using the PureLink RNA Mini Kit (Invitrogen, Thermo Fisher Scientific, 12183018A) and on-column PureLink™ DNase Set (Invitrogen, Thermo Fisher Scientific, 12185010) following the manufacturer’s instructions. Then, 1 µg of RNA was transcribed into cDNA using the High Capacity cDNA Reverse Transcription Kit with random primers (Applied Biosystems, Thermo Fisher Scientific, 4368814) and diluted to 5 ng/µl in water. RT-qPCR was performed on a CFX Opus 384 Real-Time PCR System (Bio-Rad Laboratories, 12011452) in Hard-Shell 384-Well PCR Plates (Bio-Rad Laboratories, HSP3801) with three technical replicates. Each well contained 5 µl Power SYBR Green PCR Master Mix (Applied Biosystems, Thermo Fisher Scientific, 4367659), 1.2 µl primer mix (forward and reverse primers, 1.25 µM each), 2.55 µl ddH2O and 1.25 µl cDNA (5 ng/µl). The plates were sealed with Microseal ‘B’ PCR Plate Sealing Film (Bio-Rad Laboratories, MSB1001) and the PCR reaction contained the following steps: 2 min at 50 °C, 10 min at 95 °C, then 44 cycles of: 15 sec at 95 °C, 1 min at 60 °C, plate read, and after that a final 15 sec of 95 °C followed by 60 °C - 95 °C for 15 sec to generate melting curves. Quantification cycles (Cq) were calculated using the Bio-Rad CFX Maestro software (Version 2.3, 5.3.022.1030). Calculation of relative mRNA expression levels was performed by first taking the average Cq value for each gene of the control group in three or four biological replicates. Then, the Cq values from each biological replicate and each sample were subtracted from these average values (delta Cq). To factor in primer efficiency, each primer efficiency was taken to the power of the delta Cq value for each data point (relative quantity). The normalising factor was calculated as the geometric mean of the relative quantity of all three reference genes [*PPIB* (Peptidylprolyl Isomerase B, also known as Cyclophilin B), *GAPDH* (Glyceraldehyde-3-Phosphate Dehydrogenase) and *B2M* (Beta-2-Microglobulin)]. In a final step, the relative quantity for each gene was divided by the normalising factor for each biological replicate to arrive at relative mRNA expression levels. PCR amplification efficiencies were determined for each primer pair from a calibration curve with the following cDNA concentrations: 5 ng/μl, 1 ng/μl, 0.2 ng/μl, 0.04 ng/μl, 0.008 ng/μl and 0.0016 ng/µl. The log10 values of these concentrations were plotted on the x-axis and the resulting Cq values on the y-axis. Then, a linear regression curve was fitted, and its slope was used to calculate the efficiency E with the equation: E = 10^(−1/slope). Data analysis was done in Microsoft Excel.

### Immunofluorescence

eGFP/eBFP2-β-catenin DLD-1 cells were infected with virus expressing ORF-vhhGFP constructs and selected for two days with puromycin, as described above. Then, ca. 200 000 cells were seeded on cover glasses in a 24 well plate and fixed the next day using 4% paraformaldehyde (Electron Microscopy Sciences, 15714-S) in PBS for 10 min at room temperature in the dark, followed by three washes with PBS for 5 min. Cells were permeabilised for 10 min with 0.2% Triton X-100/PBS (BioShop Canada Inc., TRX777) and then blocked for at least 30 min in blocking solution [1% Bovine Serum Albumine (BioShop Canada Inc., ALB001), 0.1% Triton X-100/PBS]. To stain for vhhGFP, samples were incubated for 1 h with MonoRab Rabbit Anti-Camelid VHH Antibody (iFluor 647, A01994-200, GenScript) diluted 1/1000 in PBS and washed again three times for 5 min with PBS. Finally, cover glasses were mounted on microscope slides with Fluoromount Aqueous Mounting Medium (Sigma-Aldrich, F4680). All steps were carried out at room temperature and samples were protected from light as much as possible. Images were acquired in the .czi format using a motorised inverted ZEISS Axio Observer microscope with the ZEN 2.5 (blue edition) software, and brightness and contrast were adjusted using Fiji (Version 1.53t).

### Inhibitor treatments

To inhibit cellular protein degradation machineries, 500 000 eGFP/eBFP2-β-catenin DLD-1 cells (uninfected or expressing ORF-vhhGFP constructs after puromycin selection) were seeded in 1 ml growth medium in a 24 well plate and treated for 18 h with 10 µM MG-132 (Cell Signaling Technology, 2194S), 1 µM TAK-243 (Selleck Chemicals, S8341), 10 µM MLN4924 (ChemieTek, CT-M4924), 1 µM Bafilomycin A (InvivoGen, tlrl-baf1) or 0.1% Dimethyl sulfoxide (DMSO, Invitrogen, Thermo Fisher Scientific, D12345).

To induce proximity between ORF-FKBP12^F36V^ constructs and eGFP-FRB*- or HaloTag-β-catenin, cells were treated with 0.5 µM Rapamycin (Selleck Chemicals, S1039), 1 µM A/C Heterodimerizer (Takara Bio USA, 635056), 0.1 µM PhosTAC7 (MedChemExpress, HY-145232), 0.2% ethanol (EtOH, house brand, 22734-P006-EAAN) or 0.2% DMSO for 24 h, 72 h or 14 days, as indicated (media and compounds were refreshed every two days).

### Incucyte analysis (cell growth and cell competition assays)

Time-lapse live-cell imaging to assess proliferation of DLD-1 cells was performed using an IncuCyte S3 Live-Cell Analysis System (Sartorius BioAnalytical Instruments) and the IncuCyte Basic Analyzer software. Regarding cell competition/coculture experiments, only signal from overexpressed fluorophores could be detected, not the fluorescent signal of endogenously expressed β-catenin fusion proteins.

For cell competition assays after CRISPR/Cas9-mediated knock-out of *CTNNB1*, eGFP/eBFP2-β-catenin DLD-1 cells expressing Cas9 were infected with pLentiguide-eGFP-P2A-Puromycin-U6-sgCTNNB1 (5’-GAAAAGCGGCTGTTAGTCAC-3’) or control pLentiguide-mCherry-P2A-Puromycin-U6-sgLacZ (5’-CCCGAATCTCTATCGTGCGG-3’) lentivirus in the presence of 8 μg/ml polybrene (Hexadimethrine bromide, Sigma, H9268). The next day, cells were treated with 1.5 µg/ml puromycin and selected for 48 h. Then, cells were lifted, counted and 100 000 cells of both sgCTNNB1/eGFP and sgLacZ/mCherry expressing cells were mixed and seeded in a clear bottom 24-well plate in 3 technical replicates in 1 ml growth medium. Remaining cells were processed for Western blot analysis to determine β-catenin knock-out efficiency. Red and green cells were quantified from images obtained using a 10× objective with 16 images per well every 6 h, over a time-course of 14 days and cells were split when they grew confluent.

For cell competition assays to validate results from the fitness screen, eGFP/eBFP2-β-catenin DLD-1 cells were infected with pLX301_NLuc_vhhGFP_P2A_TagRFP, pLX301_CSNK1D_vhhGFP_P2A_mNeonGreen or pLX301_SIAH2_vhhGFP_P2A_mNeonGreen lentivirus in the presence of 8 μg/ml Polybrene (Hexadimethrine bromide, Sigma, H9268). vhhGFP does not bind mNeonGreen or TagRFP. The next day, cells were treated with 1.5 µg/ml puromycin and selected for 48 h. Then, cells were lifted, counted and 50 000 cells of both NLuc-vhhGFP/TagRFP and CSNK1D-vhhGFP/mNeonGreen or SIAH2-vhhGFP/mNeonGreen expressing cells were mixed and seeded in a clear bottom 12-well plate. Red and green cells were quantified from images obtained using a 20× objective with 25 images per well every 12 h, and cells were split when they grew confluent.

For proliferation assays to test Rapamycin or A/C heterodimeriser induced degradation of eGFP-FRB*-β-catenin, 5 000 cells per well were seeded in 300 μl culture medium with compounds in transparent, flat-bottom 96-well plates with three technical replicates and were imaged every 12 h using a 10× objective over the course of 14 days. Confluence of cells was quantified from five images per well, and compounds and media were refreshed every 2 days.

### Statistical analysis

Statistical analysis and graphical representation was performed using GraphPad Prism [Version 9.5.1 (733)] and R (R Version 4.2.2, R Studio version 2024.12.1 Build 563).

### HiBiT-ALP5-CTNNB1 degradation assay

The anti-ALFA-tag nanobody NbALFA binds the ALP5 peptide with a Kd of 5 nM. (ALP5 amino acid sequence: PSGRLEEELRRRLSP, Steffen Frey, NanoTag Biotechnologies, personal communication, and (Götzke et al., 2019; Kilisch et al., 2021).

For experimental analysis, 1×10^6 HEK293 Flp-In T-REx HiBiT-ALP5-CTNNB1 cells were seeded in 2 ml growth medium per well in 6 well plates and induced with 1 µg/ml tetracycline (Bioshop Canada, TET701). One well was left untreated as control. The next day, cells were transfected with 1 µg pcDNA3.1-[ORF]-GSlinker-NbALFA-GSlinker-3xMyc-tag-SV40-TagRFP or TagRFP-GSlinker-3xFLAG-V5-tag only plasmid as control using 3 µl Lipofectamine 2000 Transfection Reagent as described above. Two days after transfection, cells were lifted and stained with Fixable Viability Dye eFluor 780 (1/1000 dilution in PBS, Invitrogen, Thermo Scientific, 65-0865-14) for 15 min in the dark on ice. Then, cells were washed and fixed using 1% paraformaldehyde (Electron Microscopy Sciences, 15714-S) in PBS for 10 min at room temperature in the dark, followed by another wash in PBS. Cells were permeabilised for 10 min with 0.2% Triton X-100/PBS (BioShop Canada Inc., TRX777) and then blocked for at least 30 min in blocking solution [1% Bovine Serum Albumine (BSA, BioShop Canada Inc., ALB001), 0.1% Triton X-100/PBS]. Both steps were carried out on ice in the dark. Then, cells were resuspended in anti-HiBit antibody (diluted 1/1 000 in PBS, Promega, N7200) and incubated at room temperature for 30 min in the dark, followed by two more washes. For incubation with the secondary antibody, cells were resuspended in PBS with anti-mouse IgG Alexa Fluor 488 antibody (1/1 000 dilution, Invitrogen, Thermo Scientific, A32723TR) and incubated on ice for 30 min in the dark. Then, cells were washed twice and resuspended in buffer for flow cytometry [Flow buffer: 1 mM EDTA (BioShop Canada, EDT001), 25 mM HEPES (Gibco, Thermo Scientific, 15630080), 1% BSA in PBS].

### GFP-SNAI1 protein abundance analysis

1×10^6 HEK293T cells were seeded in 2 ml growth medium per well in 6 well plates. The next day, cells were transfected with 50 ng pcDNA3.1-eGFP-SNAI1 and 1 µg pcDNA3.1-[ORF]-vhhGFP-SV40-TagRFP expressing plasmids using Lipofectamine 2000 Transfection Reagent according to the manufacturer’s recommendations. In brief, per well, plasmid DNA was diluted in 100 µl Opti-MEM (Gibco, Thermo Fisher Scientific, 11058021) and 3 µl Lipofectamine 2000 were diluted in 100 µl Opti-MEM. After 5 min incubation at room temperature, both solutions were combined, mixed gently and incubated for another 20 min at room temperature before being added drop-wise to the cells. 24 h later, eGFP expression as proxy for SNAI1 abundance was analysed using flow cytometry. TagRFP expression was used to determine transfected cells and a TagRFP-GSlinker-3xFLAG-V5-tag only plasmid was used as control for gating.

### Flow cytometry

For flow cytometry of fluorescent eGFP/eBFP2-β-catenin DLD-1 cells, samples were trypsinised, collected, washed twice with PBS and resuspended in flow buffer [1 mM EDTA (BioShop Canada, EDT001), 25 mM HEPES (Gibco, Thermo Scientific, 15630080), 1% Bovine Serum Albumine (BSA, BioShop Canada, ALB001) in PBS] containing 1 µg/ml propidium iodide (BioShop Canada, PPI888.10) as indicator of cell viability.

For flow cytometry of HEK293 or HEK293T cells transfected with TagRFP expressing plasmids, samples were trypsinised, collected, and stained with Fixable Viability Dye eFluor 780 (1/1000 dilution in PBS, Invitrogen, Thermo Scientific, 65-0865-14) for 15 min in the dark on ice before being washed twice with PBS and resuspension in flow buffer.

Live or fixed cells were kept in the dark on ice and run on a CytoFLEX LX Flow Cytometer (Beckman Coulter) with the CytExpert Software (Version 2.6.0.105, Beckman Coulter) and analysed using FlowJo Software (Version 10.7.1, BD Life Sciences). Samples were gated for cells, singlets, live cells and expression of TagRFP, if applicable. Then, β-catenin or SNAI1 abundance was determined using the median intensity of eBFP2 or eGFP signals (given in arbitrary unit, a.u.).

### Western Blot

Protein expression levels were analysed using Western blot. In brief, cells were washed twice in Dulbecco’s Phosphate Buffered Saline (PBS, Wisent Bioproducts, 311425 CL) and lysed in 8 M Urea (Invitrogen, Thermo Scientific, 15505035) or eukaryotic lysis buffer for phospho-proteins [50 mM TrisHCl (BioShop Canada, TRS001), pH 7.5, 150 mM NaCl (BioShop Canada, SOD001), 2 mM EDTA (BioShop Canada, EDT001), 10 % Glycerol (BioShop Canada, GLY002), 1% Triton X-100 (BioShop Canada, TRX777), 1% Protease Inhibitor Cocktail (Sigma-Aldrich, P2714), 20 µM Sodium Fluoride (Sigma-Aldrich, 201154), 4 mM Sodium Orthovanadate (New England Biolabs, P0758S)]. Lysates were incubated at 4 °C for at least 20 min and the spun down in a tabletop centrifuge at max speed and 4 °C to pellet and separate cell debris. Protein concentration was determined using the Pierce BCA Protein Assay (Thermo Scientific, 23225) according to the manufacturer’s instructions and a Synergy Neo microplate reader (BioTek Instruments) with the Gen5 software (Version 2.06). Samples were diluted to the desired concentration with ddH2O and denatured by adding 5x Lämmli buffer [1x formulation: 10% Glycerol, 2% SDS (BioShop Canada, SDS001), 0.02% Bromphenol Blue (Sigma, B5525), 10 mM TCEP (Aldrich, C4706), 100 mM DTT (BioShop Canada, DTT001), 62.5 mM TrisHCl, pH 6.8, in ddH2O] and boiled for 5 min at 95 °C. Per sample and well, between 2 µg and 10 µg protein was loaded on 4–15% Mini-PROTEAN TGX Stain-Free Protein Gels (Bio-Rad Laboratories, 4568086) or 26 well NuPAGE Bis-Tris Midi Protein Gels, 4 to 12% (Invitrogen, Thermo Scientific, WG1403), with Precision Plus Protein All Blue Standards (Bio-Rad Laboratories, 1610373) as reference. SDS-PAGE of Mini-PROTEAN gels was run with 1x Tris/Gylcine buffer [192 mM Glycine (BioShop Canada, GLN001), 25 mM TRIS, 0.1 % SDS, in ddH2O] and SDS-PAGE of NuPAGE Bis-Tris gels was run with 1x MOPS SDS buffer [50 mM MOPS (BioShop Canada, MOP005), 50 mM Tris Base, 0.1% SDS, 1 mM EDTA] and proteins were transferred to nitrocellulose membranes (0.45 μm, Thermo Scientific, 88018) with 1x Transfer Buffer [10% Methanol (BioShop Canada, MET302), 95.9 mM Glycine, 12 mM TRIS in ddH2O]. Successful transfer was confirmed with Ponceau S solution (Sigma, P7170) and membranes were blocked in 5% Skim Milk/TBST (BioShop Canada, SKI400) or 5% BSA/TBST (BioShop Canada, ALB001, for Phospho-β-catenin blots) at room temperature for at least 1 h. Then, membranes were incubated with primary antibodies overnight at 4 °C. The next day, membranes were washed at least three times for 7 min in TBST [50 mM TrisHCl, pH 7.5, 150 mM NaCl (BioShop Canada, SOD001), 0.1% Tween-20 (BioShop Canada, TWN510)], and then incubated in secondary antibodies for 1 h at room temperature, followed by another three washes as above before imaging using a ChemiDoc MP Imaging System (Bio-Rad Laboratories) and SuperSignal West Pico PLUS Chemiluminescent Substrate (Thermo Scientific, 34580) or SuperSignal West Femto Maximum Sensitivity Substrate (Thermo Scientific, 34096). Primary antibodies with HRP-conjugation were incubated at room temperature for 30 min before washing and imaging. If necessary, HRP signal was quenched using 10 % Acetic Acid (Caledon Laboratories, 1000-1-29) at room temperature for 20 min, followed by three washes for 20 min in TBST. Western blot quantification was performed by densitometry of β-catenin normalised to GAPDH signal after background subtraction in Fiji (Version 1.53t).

### Immunoprecipitation

Proteins were isolated from 6×10^6 eGFP/eBFP2-β-catenin DLD-1 cells using eukaryotic lysis buffer [50 mM TrisHCl (BioShop Canada, TRS001), pH 7.5, 150 mM NaCl (BioShop Canada, SOD001), 2 mM EDTA (BioShop Canada, EDT001), 10 % Glycerol (BioShop Canada, GLY002), 1% Triton X-100 (BioShop Canada, TRX777), 1% Protease Inhibitor Cocktail (Sigma-Aldrich, P2714)] by incubating at 4 °C for at least 20 min followed by spinning down in a tabletop centrifuge at max speed (4 °C) and removal of the cell debris pellet. Then, Input control samples were taken and prepared for Western blot. The remaining sample was divided into 2 tubes and incubated with 3 µl of anti-β-catenin antibody (Cell Signaling Technology, #9587) or equivalent amount of Rabbit IgG Isotype Control (Invitrogen, Thermo Fisher Scientific, 31235) for 1 h at 4 °C. In the meantime, 50 µl Pierce Protein A Agarose slurry (Thermo Scientific, 20333) per sample were washed 3x in eukaryotic lysis buffer (centrifuged in between at 12 000 xg for 30 sec) and then blocked in 5% BSA (BioShop Canada Inc., ALB001)/TBST [50 mM TrisHCl, pH 7.5, 150 mM NaCl (BioShop Canada, SOD001), 0.1% Tween-20 (BioShop Canada, TWN510)] for 1 h in the cold room, followed again by 3 washes as previously. Then, the washed beads were distributed evenly among the samples, followed by another 3 h incubation in the cold room. Then, the beads were collected by centrifugation (12 000 xg, 30 sec) and supernatant was removed. Samples were washed twice for 5 min at 4 °C with each of the following wash buffers: 1^st^ wash buffer: eukaryotic lysis buffer; 2^nd^ wash buffer (high salt): eukaryotic lysis buffer with 500 mM NaCl; 3^rd^ wash buffer (low salt): eukaryotic lysis buffer without NaCl]. After the last wash, beads were collected by centrifugation as previously, supernatant was removed as completely as possible and beads were resuspended in 40 µl 1x Lämmli buffer [10% Glycerol, 2% SDS (BioShop Canada, SDS001), 0.02% Bromphenol Blue (Sigma, B5525), 10 mM TCEP (Aldrich, C4706), 100 mM DTT (BioShop Canada, DTT001), 62.5 mM TrisHCl, pH 6.8, in ddH2O] and boiled for 5 min at 95 °C, before being analysed by Western blot.

### Induced-proximity screens

Pooled ORFeome library generation and lentivirus production are described in Poirson et al., 2024. ORFeome-vhhGFP libraries were packaged into lentiviral particles. The endogenously tagged colorectal cancer cell line DLD-1 CTNNB1^eBFP2/eGFP^ (clone #28) was transduced in the presence of 8 µg/ml polybrene (Hexadimethrine bromide, Sigma, H9268) at low multiplicity of infection so that approximately 50% cells survived after puromycin (1.5 μg/ml) selection and untransduced cells were fully eliminated.

For the induced-proximity Fluorescence Activated Cell Sorting (FACS) screen for β-catenin protein abundance, cells were expanded for three days after puromycin selection, then washed in PBS, trypsinised, collected in growth medium, washed in PBS again and resuspended in sorting buffer [1 mM EDTA (BioShop Canada, EDT001), 25 mM HEPES (Gibco, Thermo Scientific, 15630080), 1% Bovine Serum Albumine (BSA, BioShop Canada, ALB001) in PBS] containing 1 µg/ml propidium iodide (BioShop Canada, PPI888.10) as indicator of cell viability. 5×10^6 unsorted cells were kept as reference for baseline. Gating of live, single, eBFP2/eGFP double positive cells was done with untransduced control cells and the respective ‘eBFP2/eGFP double negative’ population was sorted in biological duplicates using a BD FACS Melody (BD Biosciences) and a BD FACSymphony S6 SE (BD Biosciences) cell sorter (both at the University of Toronto Centre for Immune Analytics Core Facility, RRID: SCR_027612). 50 000 and 175 000 cells were sorted as double negative populations for genomic DNA isolation.

For the induced-proximity dropout screen for cell fitness, > 250-fold coverage of the library was maintained throughout the experiment. After Puromycin selection, 5×10^6 cells per replicate were collected as T0 (baseline) samples and 5×10^6 of the remaining cells per replicate in two replicates were continued to culture and split when necessary. As control, untagged DLD-1 cells were transduced with the library and cultured in the same way. 14 days after puromycin selection, 5×10^6 cell per replicate were collected for genomic DNA isolation (T1).

Genomic DNA was directly extracted using the Qiagen DNeasy Blood and Tissue kit (Qiagen, 69504) for both screens and ORFeome next-generation sequencing was performed as described in Poirson et al., 2024. The samples were submitted to the Donnelly Sequencing Center at the University of Toronto (https://thedonnellycentre.utoronto.ca/donnelly-sequencing-centre) for sequencing.

### Analysis of ORFeome sequencing data and GO-term enrichment

FASTQ files from Illumina sequencing runs for the ORFeome screen were mapped to an index of the ORFeome reference sequences using Kallisto (Version 0.44.0, (Bray et al., 2016). Log_2_(fold change) and adjusted p-values of differentially expressed count data was calculated using DESeq2 (Version 1.46.0, (Love et al., 2014). For the FACS screen, a 5% cut-off for adjusted p-value and 4-fold change in normalised read counts between the sorted (eBFP2/eGFP double negative) population and the unsorted population was used. For the Fitness screen, a 5% cut-off for adjusted p-value and 2-fold change in normalised read counts between T1 and T0 was used. Enrichment of GO molecular functions terms was performed using g:Profiler (https://biit.cs.ut.ee/gprofiler/gost, (Reimand et al., 2016).

### RNA-sequencing

For bulk RNA sequencing, DLD-1 CTNNB1^eGFP-FRB*/eGFP-FRB*^ cells (clone #43) were infected with lentivirus expressing CSNK1D-FKBP12^F36V^ and selected for 2 days with puromycin (1.5 µg/ml). Then, 500 000 cells were seeded in a 12 well plate and treated with 1 µM A/C Heterodimeriser (Takara Bio USA, Inc., 635056) or 0.2 % EtOH as vehicle control for 8 h or 16 h. RNA was isolated using the PureLink RNA Mini Kit (Invitrogen, Thermo Fisher Scientific, 12183018A) and on-column PureLink™ DNase Set (Invitrogen, Thermo Fisher Scientific, 12185010) following the manufacturer’s instructions as described above. The samples were submitted to the Donnelly Sequencing Center at the University of Toronto (https://thedonnellycentre.utoronto.ca/donnelly-sequencing-centre) for sequencing. The quality of RNA was measured using a BioAnalyzer (Agilent) and RNAseq libraries for 3 biological replicates of each sample were derived using NEBNext Poly(A) mRNA Magnetic isolation module followed by NEBNext Ultra II Directional RNA library prep kit. Libraries were run on a Novaseq6000 SP300c kit (Illumina), generating 150 bp paired-end FASTQ files. The trimmed reads were aligned to the human genome assembly GRCh38 (Genome Reference Consortium) using Kallisto (Version 0.44.0, (Bray et al., 2016), following default parameters. Principal component analysis and differential expression analysis were performed using the DESeq2 package with R (R Version 4.2.2, R Studio version 2024.12.1 Build 563, https://www.r-project.org/) using default parameters (DESeq2 Version 1.46.0, (Love et al., 2014). Enrichment of gene ontology (GO) biological processes (BP) terms was performed using g:Profiler (https://biit.cs.ut.ee/gprofiler/gost). Driver terms derived from the most downregulated genes (5% cut-off for adjusted p-value and 2-fold change in normalised read counts between A/C and EtOH) with a maximum term size of 350 are shown (Reimand et al., 2016). Hallmark Gene Set Enrichment Analysis (GSEA) were performed on a ranked list of genes using the GSEA-Preranked tool within the GSEA software (version 4.4.0) and standard parameters (Subramanian et al., 2005).

### Genome-wide CRISPR/Cas9 Knock out (KO) screen

The CRISPR/Cas9 KO screen was performed in DLD-1 CTNNB1^eGFP-FRB*/eGFP-FRB*^ cells (clone #43) after stable, lentiviral integration of CSNK1D-FKBP12^F36V^-HAtag_T2A_Hygromycin and hygromycin selection using the 70K TKOv3 library (Addgene Pooled Library #90294)(Hart et al., 2015). Around 200 x10^6 cells were infected with TKOv3 virus at a multiplicity of infection (MOI) of 0.3 in the presence of 8 µg/ml polybrene in 30x 15 cm plates with 25 ml growth medium each. The next day, media was removed and puromycin selection (1.5 µg/ml) started and continued for 3 days, cells were lifted after 2 days to increase selection pressure. Then, cells were harvested, counted and 35×10^6 cells were collected as T0 sample and stored at −80°C for later processing. 25 x10^6 of the remaining cells were seeded for each replicate (3x treatment and 3x control) and treated with 1 µM A/C Heterodimeriser (Takara Bio USA, Inc., 635056) or 0.2 % EtOH as vehicle control. Media was changed every 48 h to refresh drugs and cells were split every 4 days. A minimum of 200-fold library coverage was maintained for all replicates throughout the screen. Cell pellets were collected at 8 days (Midpoint) and 16 days (Endpoint) after T0 and stored at −80°C for later processing. Genomic DNA extraction from frozen cell pellets was performed using the QIAamp DNA Blood Maxi Kit (Qiagen, 51194) and sgRNA sequences were amplified and barcoded with Illumina TruSeq adapters. The barcoded PCR products were then gel purified and sequenced on a NextSeq2000 instrument (Illumina) at a read depth of 200-fold for all samples. The samples were submitted to the Donnelly Sequencing Center at the University of Toronto (https://thedonnellycentre.utoronto.ca/donnelly-sequencing-centre) for sequencing. Analysis of screen sequences was performed following the MAGeCK (version 0.5.9.2) and DrugZ (version 1.1.0.2) pipelines (Colic et al., 2019; Li et al., 2014).

**Table 1:**
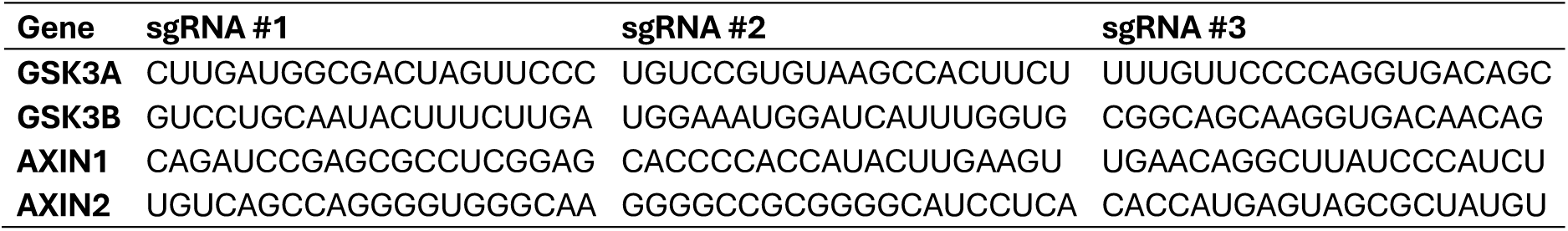
sgRNA sequences used for gene knock-out with RNPs.

**Table 2:**
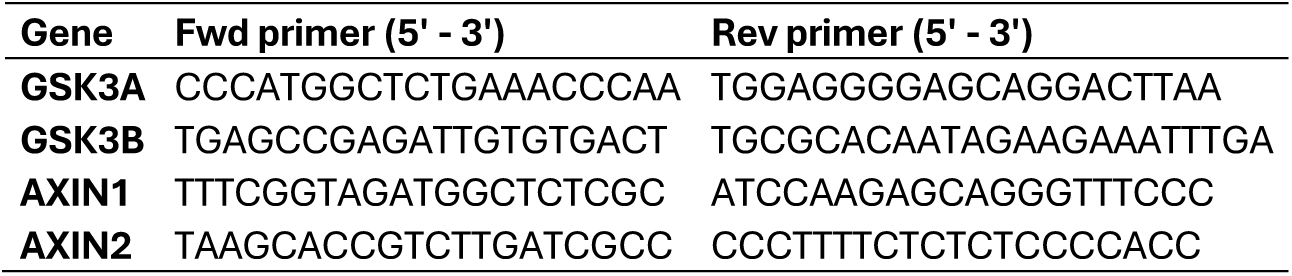
Primer sequences used to amplify regions targeted by sgRNAs.

**Table 3:**
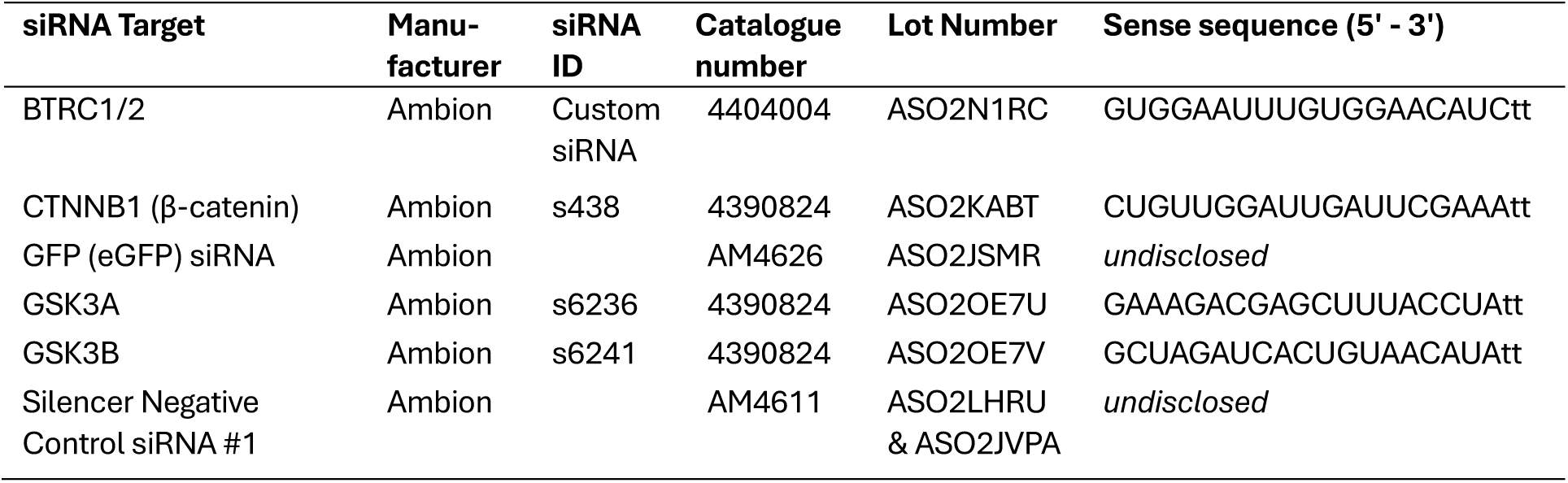
siRNAs used in this study.

**Table 4:**
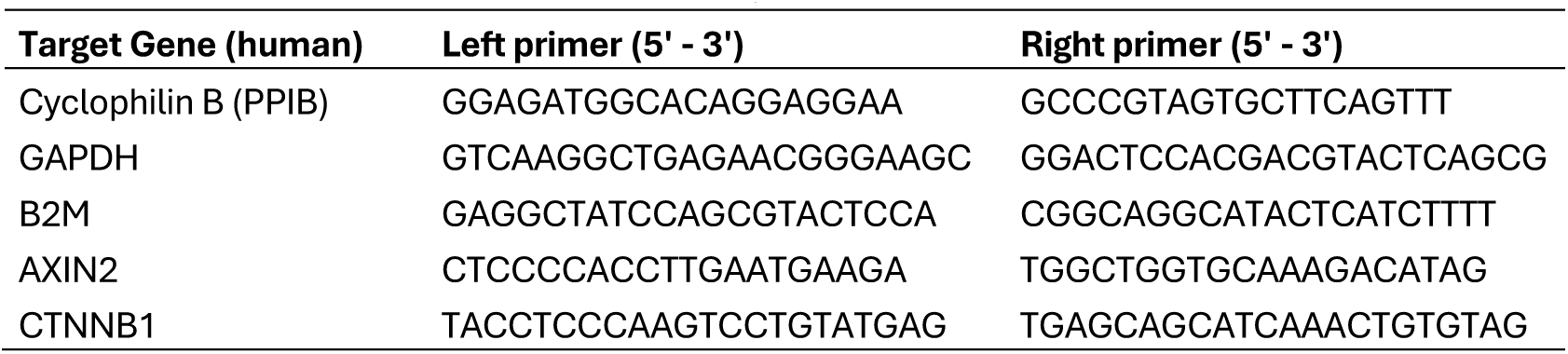
Primers for RT-qPCR used in this study.

**Table 5:**
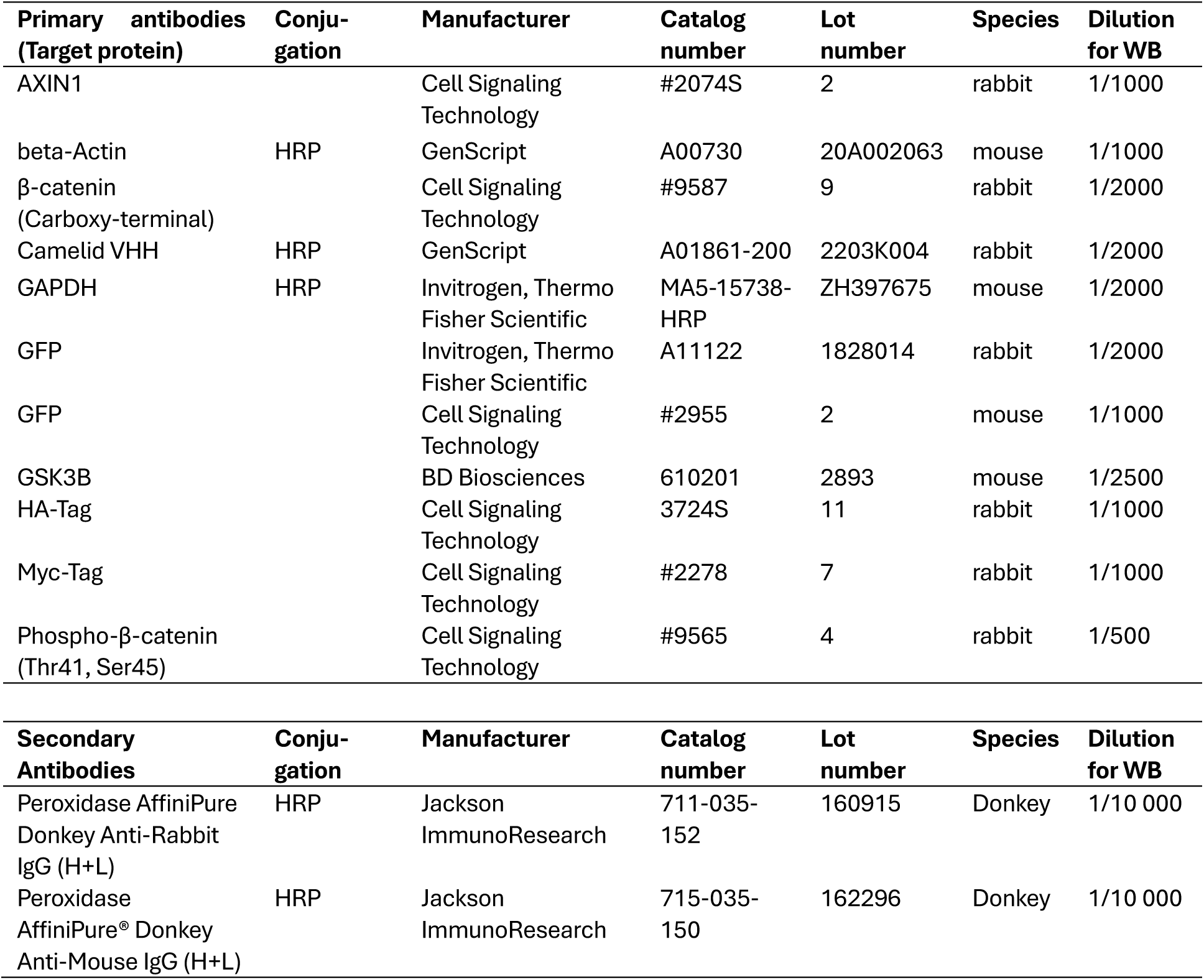
Antibodies used for Western blot (WB) in this study.

## ACKNOWLEDGEMENTS

L.W. was supported by the Deutsche Forschungsgemeinschaft (DFG, German Research Foundation, project no. 533617596) and the Charles H. Best Postdoctoral Fellowship (2023). This work was also supported by a Canadian Institutes of Health Research grant (PJT-175160) to SA. The authors would like to thank the Donnelly Sequencing Center at the University of Toronto (https://thedonnellycentre.utoronto.ca/donnelly-sequencing-centre) for next-generation sequencing services, the University of Toronto Centre for Immune Analytics Core Facility (RRID: SCR_027612) for cell sorting services and the Centre for Pharmaceutical Oncology at the University of Toronto for providing access to equipment and resources. We would like to thank all members of the Angers and Taipale labs for valuable discussion and feedback.

## DATA AVAILABILITY

RNA-Seq data generated from this study is deposited in GEO (GSE324133).

## AUTHOR CONTRIBUTIONS

Conceptualization: L.W, S.A.; Methodology: L.W., J.P., G.M., S.L., Y.K., M.A. Formal analysis: L.W., J.P., G.M.; Investigation: L.W.; Writing - original draft: L.W, S.A.; Writing - review & editing: L.W., J.P., G.M., S.L., Y.K., M.A., M.T. & S.A.; Supervision: M.T. & S.A.; Funding acquisition: L.W., M.T. & S.A.

