## Supplement for "Phosphorylation-driven Targeted Protein Degradation of Oncogenic β-catenin"

- **Supplementary Figures S1 – S5**
- **Constructs for endogenous tagging of CTNNB1 (5' – 3')**
- **Constructs for stable integration into HEK293 Flp-In T-REx cells (5' – 3')**

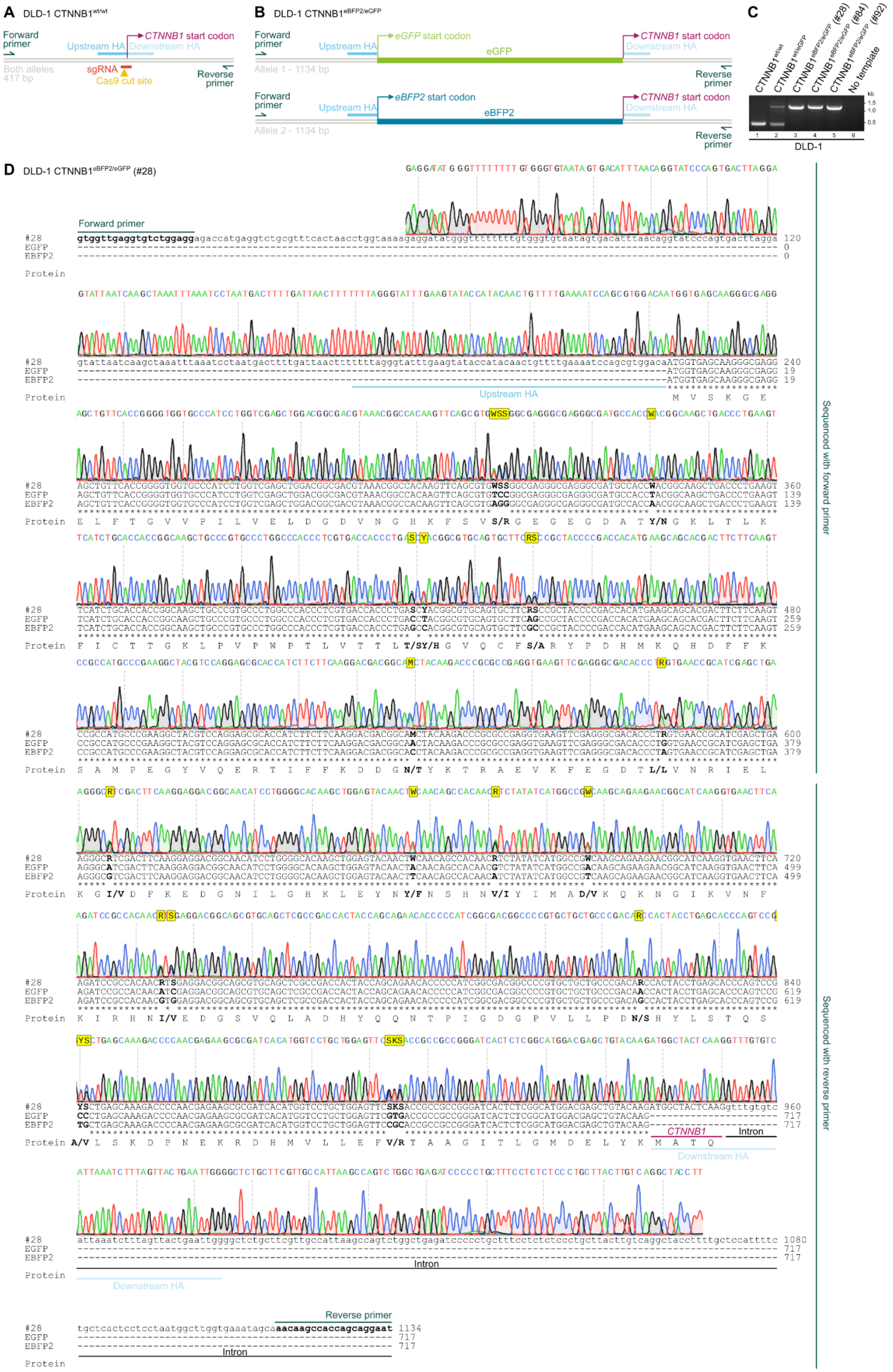
A DLD-1 CTNNB1wt/wt
Forward primer
Upstream HA
CTNNB1 start codon
Downstream HA
Both alleles 417 bp
sgRNA
Cas9 cut site
Reverse primer
B DLD-1 CTNNB1eBFP2/eGFP
eGFP start codon
eGFP
Allele 1 - 1134 bp
eBFP2 start codon
eBFP2
Allele 2 - 1134 bp
C
CTNNB1wt/wt
CTNNB1wt/eGFP
CTNNB1eBFP2/eGFP (#28)
CTNNB1eBFP2/eGFP (#84)
CTNNB1eBFP2/eGFP (#92)
No template
kb
1.5
1.0
0.5
DLD-1
D DLD-1 CTNNB1eBFP2/eGFP (#28)
#28
EGFP
EBFP2
Protein
Intron
Sequenced with forward primer
Sequenced with reverse primer

**Suppl. Fig. 1: Endogenous tagging of β-catenin with eBFP2 and eGFP in the colorectal cancer cell line DLD-1**
**A, B)** Schematic depiction of the PCR products obtained from amplifying the translational start site of the endogenous *CTNNB1* (the gene encoding β-catenin) locus in untagged (wild-type, wt) or double tagged (eBFP2/eGFP) DLD-1 cells. Indicated are binding sites of the single guide RNA (sgRNA) and homology arms (HA) used for CRISPR/Cas9-mediated endogenous tagging of *CTNNB1*, with eGFP (Enhanced Green Fluorescent Protein) and eBFP2 (Enhanced Blue Fluorescent Protein 2). bp = base pairs
**C)** PCR products amplified from the endogenous *CTNNB1* locus in untagged (wt), single tagged (wt/eGFP), or double tagged (eBFP2/eGFP) DLD-1 cells. Three different double tagged single cell clones are shown (clones #28, #84, and #92). kb = kilo base pairs
**D)** Sanger sequencing of the PCR product obtained from amplifying the translational start site of the double tagged DLD-1 cell clone #28 aligned to the nucleotide sequences of eGFP, eBFP2, or the predicted PCR amplicon sequence shows a perfect integration of both eGFP and eBFP2 at the 5' end of *CTNNB1*, corresponding to an N-terminal protein tag. The sequences of eGFP and eBFP2 are very similar, with only 22 bp or 14 amino acids differences between the fluorophores.

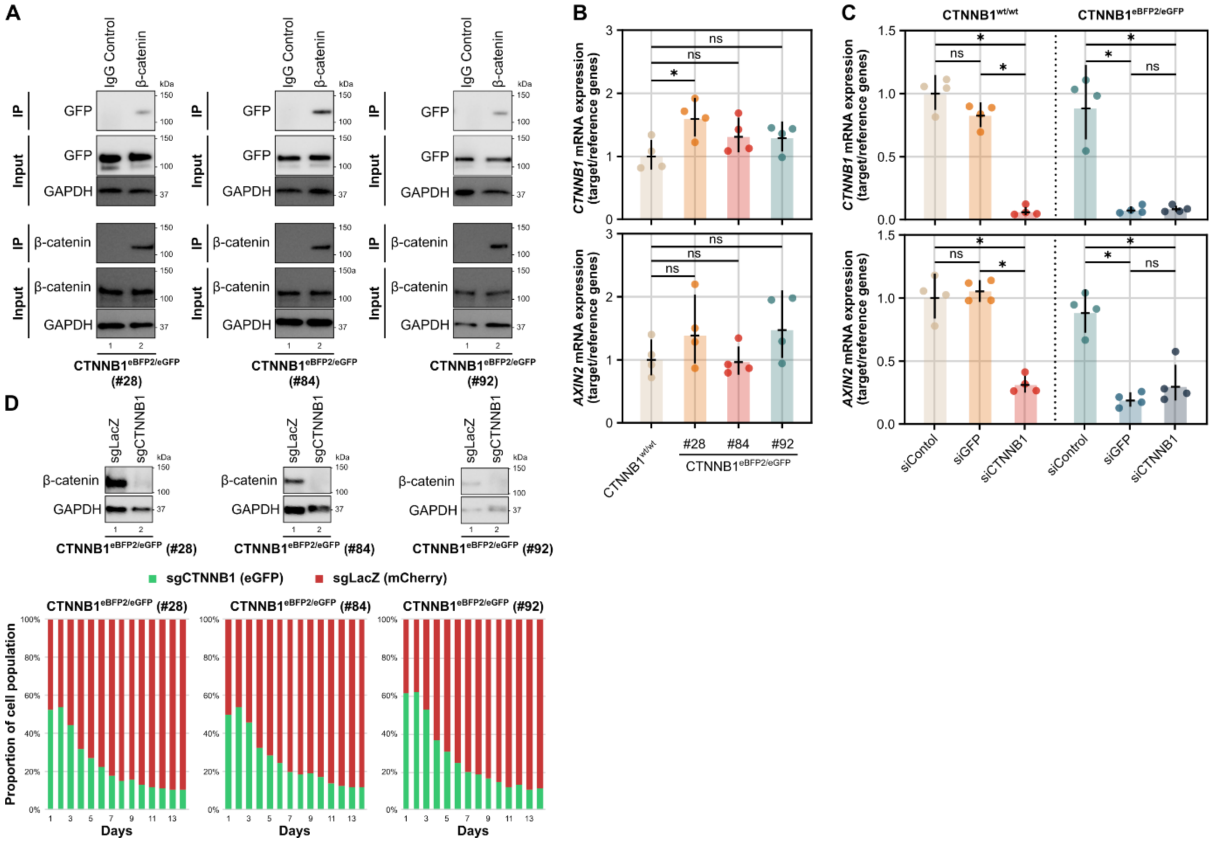


**Suppl. Fig. 2: Double tagged eBFP2/eGFP-β-catenin is functional in DLD-1 cells**

**A)** Immunoprecipitation (IP) experiments with anti-β-catenin antibody confirmed interaction between β-catenin and eGFP in double tagged DLD-1 CTNNB1$^{eBFP2/eGFP}$ single cell clones #28, #84, and #92. kDa = Kilodalton, GAPDH served as loading control.

**B)** *CTNNB1* and *AXIN2* (a universal Wnt/β-catenin target gene) mRNA expression remained mostly unchanged between untagged and double tagged DLD-1 CTNNB1$^{eBFP2/eGFP}$ single cell clones #28, #84, and #92, as determined by reverse-transcription quantitative polymerase chain reaction (RT-qPCR). Fold change data was normalised to untagged cells and *GAPDH*, *B2M* and *PPIA* served as reference genes. Individual data points from four independent experiments are shown with geometric mean and geometric standard deviation. Statistical significance was determined by one-way ANOVA followed by Dunnett's multiple comparisons test. * Padj ≤ 0.05, ns = not significant.

**C)** Knock-down of GFP regulated *CTNNB1* and *AXIN2* in double tagged DLD-1 CTNNB1$^{eBFP2/eGFP}$ cells. Untagged or double tagged DLD-1 CTNNB1$^{eBFP2/eGFP}$ cells (clone #28) were transfected with the indicated siRNAs and harvested for RT-qPCR analysis 72 h later. siGFP targeted both eGFP and eBFP2 due to sequence similarities. Fold change data was normalised to untagged cells and *GAPDH*, *B2M* and *PPIA* served as reference genes. Individual data points from four independent experiments are shown with geometric mean and geometric standard deviation. Statistical significance was determined by one-way ANOVA followed by Tukey's multiple comparisons test. * Padj ≤ 0.0001, ns = not significant

**D)** β-catenin knock-out impaired proliferation of double-tagged DLD-1 cells. DLD-1 CTNNB1$^{eBFP2/eGFP}$ single cell clones #28, #84, and #92 were infected with virus encoding eGFP and single guide RNA targeting CTNNB1 (sgCTNNB1) for CRISPR/Cas9-mediated knock-out. Control cells were infected with virus encoding mCherry and control sgLacZ. Reduced β-catenin abundance in selected cell pools after sgCTNNB1 expression was confirmed by Western blot, kDa = Kilodalton, GAPDH served as loading control. Then, sgCTNNB1 (eGFP)- and sgLacZ (mCherry)-expressing cells were mixed and imaged using a time-lapse microscope (Incucyte) over the course of 14 days. In this co-culture experiment, cells with β-catenin knock-out proliferated less than their β-catenin expressing counterparts and dropped out from the population over time. Importantly, the expression of endogenous eGFP-β-catenin was below the detection limit of the Incucyte time-lapse microscope and only the signal of overexpressed eGFP was strong enough to be detected.

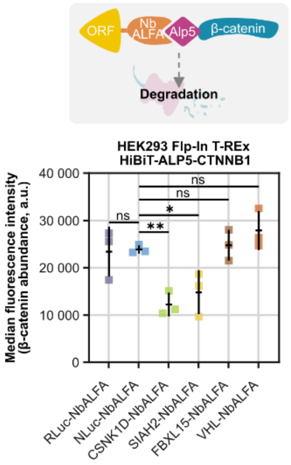


**Suppl. Fig. 3: β-catenin is degraded by CSNK1D and SIAH2 after proximity is induced by the NbALFA-ALP5-system**

HEK293 Flp-In T-REx cells stably expressing HiBiT-ALP5-β-catenin under a TET-inducible promoter were transfected with the indicated -NbALFA fusion proteins and HiBiT expression was measured using flow cytometry as read-out for β-catenin abundance. NbALFA binds to ALP5, a derivate of the ALFA-tag. Individual data points of three independent experiments are shown with mean and standard deviation. A minimum of 9,900 cells per condition were recorded in the final gate. Statistical significance was assessed by one-way ANOVA followed by Dunnett's multiple comparisons test comparing all -NbALFA groups to NLuc-NbALFA. * Padj ≤ 0.05, ** Padj ≤ 0.01 RLuc - Renilla luciferase; NLuc - NanoLuc luciferase; NbALFA - nanobody recognising ALFA-tag; a.u. - arbitrary unit

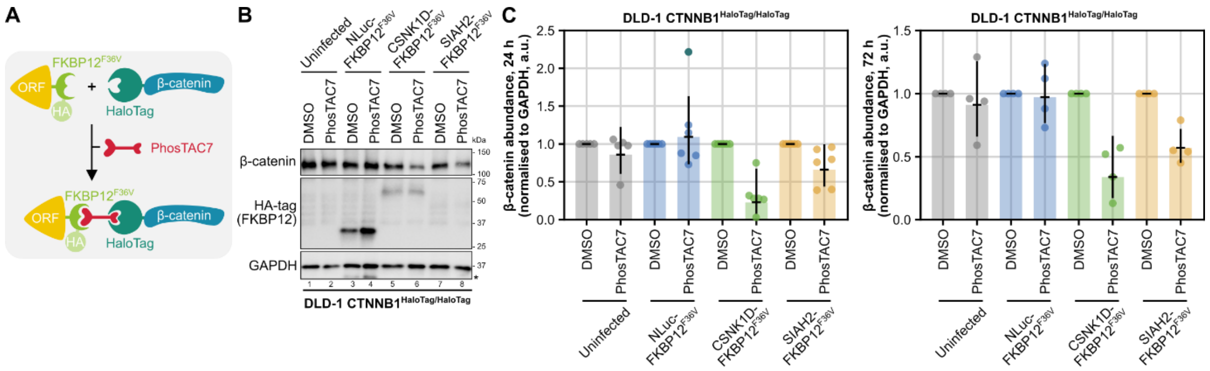


**Suppl. Fig. 4: PhosTAC7-induced proximity to CSNK1D degrades β-catenin**

**A)** Schematic depiction of induced proximity between HaloTag-β-catenin and HA- and FKBP12$^{F36V}$- tagged Open Reading Frames (ORFs) using the heterobifunctional molecule PhosTAC7.

**B, C)** PhosTAC7 induced dimerisation of β-catenin and CSNK1D induces β-catenin degradation. Double tagged DLD-1 CTNNB1$^{HaloTag/HaloTag}$ cells (clone #97) were infected with virus expressing the indicated -FKBP12$^{F36V}$ fusion proteins, selected with puromycin for 2 days and then treated with PhosTAC7 (0.1 µM) or equivalent volumes of DMSO for 24 h or 72 h. **B)** representative Western Blot of n = 4 biological replicates after 72 h treatment; * signal from previous staining; kDa = Kilodalton; GAPDH served as loading control. **C)** β-catenin abundance was quantified from Western Blots after 24 h or 72 h of treatment and normalised to GAPDH and DMSO treatment. Fold change data of n = 5 or 6 biological replicates (24 h) and n = 4 biological replicates (72 h) with geometrical mean and geometric standard deviation is shown. a.u. - arbitrary unit

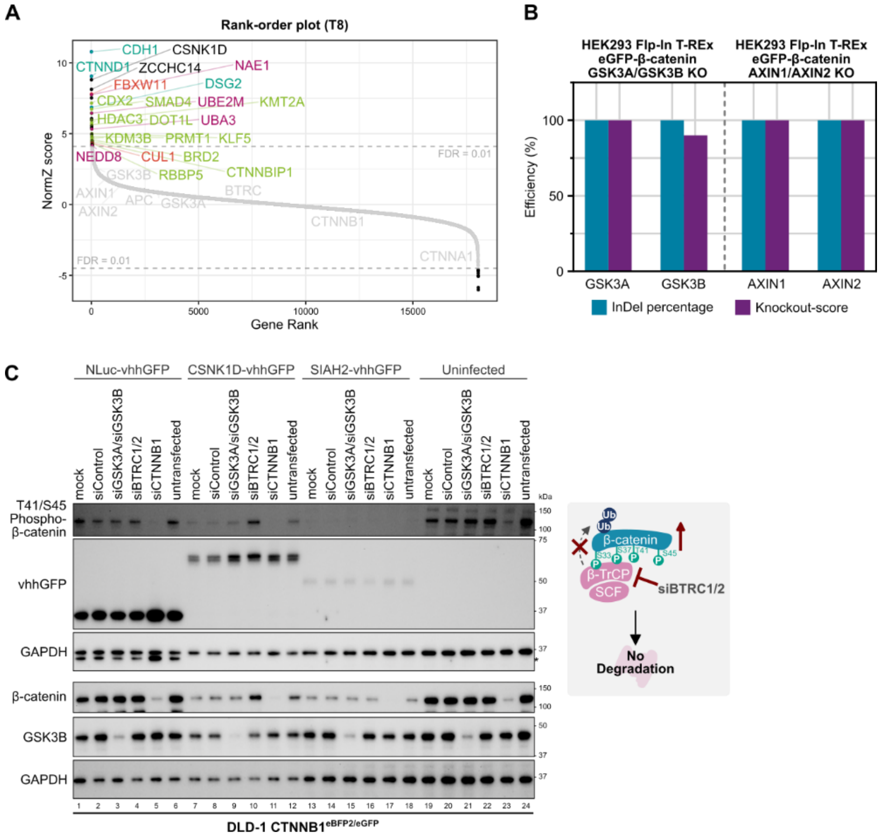


**Suppl. Fig. 5: β-TrCP2 (BTRC2/FBXW11) is required for CSNK1D-dependent degradation of β-catenin**

**A)** Rank order plot depicting the gene level summary of sgRNAs that regulate CSNK1D-dependent β-catenin degradation identified in the genome-wide CRISPR/Cas9 positive selection screen at Midpoint (T8). Black dots represent regulators with FDR < 0.01. Highlighted are genes involved in cell-cell contacts (dark green), transcriptional regulation (light green), neddylation (pink), the SCF$^{\beta\text{-TrCP}}$ E3 ubiquitin ligase complex (red, FBXW11 is also known as BTRC2), or others (black).

**B)** ICE (Inference of CRISPR Edits, https://ice.editco.bio) analysis of GSK3A/GSK3B or AXIN1/AXIN2 double knock-out (KO) HEK293 Flp-In T-REx GFP-CTNNB1 cell lines after CRISPR ribonucleoprotein (RNP) transfection confirms target gene knock-out.

**C)** CSNK1D-dependent degradation of β-catenin is BTRC dependent. The human genes BTRC1 and BTRC2 encode beta-TrCP1 and beta-TrCP2 (also known as FBXW11), which are part of the E3 ubiquitin ligase complex that ubiquitinylates β-catenin in the context of the destruction complex. SiRNA-mediated knock-down of BTRC1/2 rescued CSNK1D-dependent degradation of β-catenin and lead to accumulation of T41/S45 phosphorylated β-catenin, whereas knock-down of GSK3A and GSK3B had no effect. DLD-1 CTNNB1$^{eBFP2/eGFP}$ cells (clone #28) were infected with virus expressing the indicated -vhhGFP fusion proteins, selected with puromycin for 2 days and then transfected with the indicated siRNAs and analysis by Western Blot 48 h later. * signal from previous staining; kDa = Kilodalton; GAPDH served as loading control.

RLuc - Renilla luciferase; NLuc - NanoLuc luciferase; vhhGFP - nanobody binding to eGFP and eBFP2

**Constructs for endogenous tagging of CTNNB1 (5’ – 3’)**

**EGFP**
atggtgagcaagggcgaggagctgttcaccggggtggtgcccatcctggtcgagctggacggcgacgtaaacggccacaagttcagcgtgtccg
gcgagggcgagggcgatgccacctacggcaagctgaccctgaagttcatctgcaccaccggcaagctgcccgtgccctggcccaccctcgtg
accaccctgacctacggcgtgcagtgcttcagccgctaccccgaccacatgaagcagcacgacttcttcaagtccgccatgcccgaaggcta
cgtccaggagcgcaccatcttcttcaaggacgacggcaactacaagacccgcgccgaggtgaagttcgagggcgacaccctggtgaaccgc
atcgagctgaagggcatcgacttcaaggaggacggcaacatcctggggcacaagctggagtacaactacaacagccacaacgtctatatcat
ggccgacaagcagaagaacggcatcaaggtgaacttcaagatccgccacaacatcgaggacggcagcgtgcagctcgccgaccactacca
gcagaacacccccatcggcgacggccccgtgctgctgcccgacaaccactacctgagcacccagtccgccctgagcaaagaccccaacg
agaagcgcgatcacatggtcctgctggagttcgtgaccgccgccgggatcactctcggcatggacgagctgtacAAG

**EBFP2**
Atggtgagcaagggcgaggagctgttcaccggggtggtgcccatcctggtcgagctggacggcgacgtaaacggccacaagttcagcgtgagg
ggcgagggcgagggcgatgccaccaacggcaagctgaccctgaagttcatctgcaccaccggcaagctgcccgtgccctggcccaccctcg
tgaccaccctgagccacggcgtgcagtgcttcgcccgctaccccgaccacatgaagcagcacgacttcttcaagtccgccatgcccgaagg
ctacgtccaggagcgcaccatcttcttcaaggacgacggcacctacaagacccgcgccgaggtgaagttcgagggcgacaccctagtgaac
cgcatcgagctgaagggcgtcgacttcaaggaggacggcaacatcctggggcacaagctggagtacaacttcaacagccacaacatctatat
catggccgtcaagcagaagaacggcatcaaggtgaacttcaagatccgccacaacgtggaggacggcagcgtgcagctcgccgaccactac
cagcagaacacccccatcggcgacggccccgtgctgctgcccgacagccactacctgagcacccagtccgtgctgagcaaagaccccaa
cgagaagcgcgatcacatggtcctgctggagttccgcaccgccgccgggatcactctcggcatggacgagctgtacAAG

**EGFP-FRB***
Atggtgagcaagggcgaggagctgttcaccggggtggtgcccatcctggtcgagctggacggcgacgtaaacggccacaagttcagcgtgtcc
ggcgagggcgagggcgatgccacctacggcaagctgaccctgaagttcatctgcaccaccggcaagctgcccgtgccctggcccaccctcgt
gaccaccctgacctacggcgtgcagtgcttcagccgctaccccgaccacatgaagcagcacgacttcttcaagtccgccatgcccgaaggct
acgtccaggagcgcaccatcttcttcaaggacgacggcaactacaagacccgcgccgaggtgaagttcgagggcgacaccctggtgaaccg
catcgagctgaagggcatcgacttcaaggaggacggcaacatcctggggcacaagctggagtacaactacaacagccacaacgtctatatca
tggccgacaagcagaagaacggcatcaaggtgaacttcaagatccgccacaacatcgaggacggcagcgtgcagctcgccgaccactacc
agcagaacacccccatcggcgacggccccgtgctgctgcccgacaaccactacctgagcacccagtccgccctgagcaaagaccccaac
gagaagcgcgatcacatggtcctgctggagttcgtgaccgccgccgggatcactctcggcatggacgagctgtacaag (EGFP)-
ggcgcgcgcgccaatcgaattaattctaga (linker) –
atcctctggcatgagatgtggcatgaaggcctggaagaggcatctcgtttgtactttggggaaaggaacgtgaaaggcatgtttgaggtgctggagc
ccttgcatgctatgatggaacggggcccccagactctgaaggaaacatcctttaatcaggcctatggtcgagatttaatggaggcccaagagtgg
tgcaggaagtacatgaaatcagggaatgtcaaggacctcctccaagcctgggacctctattatcatgtgttccgacgaatctcaaag (FRB*) –
gtgtacAAG (linker)

**HaloTag-HA-tag**
atggcagaaatcggtactggctttccattcgacccccattatgtggaagtcctgggcgagcgcatgcactacgtcgatgttggtccgcgcgatggc
acccctgtgctgttcctgcacggtaacccgacctcctcctacgtgtggcgcaacatcatcccgcatgttgcaccgacccatcgctgcattgctc
cagacctgatcggtatgggcaaatccgacaaaccagacctgggttatttcttcgacgaccacgtccgcttcatggatgccttcatcgaagccctg
ggtctggaagaggtcgtcctggtcattcacgactggggctccgctctgggtttccactgggccaagcgcaatccagagcgcgtcaaaggtattgc
atttatggagttcatccgccctatcccgacctgggacgaatggccagaatttgcccgcgagaccttccaggccttccgcaccaccgacgtcggc
cgcaagctgatcatcgatcagaacgtttttatcgagggtacgctgccgatgggtgtcgtccgcccgctgactgaagtcgagatggaccattaccg
cgagccgttcctgaatcctgttgaccgcgagccactgtggcgcttcccaaacgagctgccaatcgccggtgagccagcgaacatcgtcgcgct
ggtcgaagaatacatggactggctgcaccagtcccctgtcccgaagctgctgttctggggcaccccaggcgttctgatcccaccggccgaagc
cgctcgcctggccaaaagcctgcctaactgcaaggctgtggacatcggcccgggtctgaatctgctgcaagaagacaacccggacctgatcg
gcagcgagatcgcgcgctggctgtcgacgctcgagatttccggc (HaloTag)-
gagccgaccact (linker)- taccccctatgatgtccccgactacgcc (HA-tag)- ctgtacaag (linker)

**Constructs for stable integration into HEK293 Flp-In T-REx cells (5’ – 3’)**

**GFP-CTNNB1 (wild type)**
Atggtgagcaagggcgaggagctgttcaccggggtggtgcccatcctggtcgagctggacggcgacgtaaacggccacaagttcagcgtgtcc
ggcgagggcgagggcgatgccacctacggcaagctgaccctgaagttcatctgcaccaccggcaagctgcccgtgccctggcccaccctcgt
gaccaccctgacctacggcgtgcagtgcttcagccgctaccccgaccacatgaagcagcacgacttcttcaagtccgccatgcccgaaggct
acgtccaggagcgcaccatcttcttcaaggacgacggcaactacaagacccgcgccgaggtgaagttcgagggcgacaccctggtgaaccg
catcgagctgaagggcatcgacttcaaggaggacggcaacatcctggggcacaagctggagtacaactacaacagccacaacgtctatatca
tggccgacaagcagaagaacggcatcaaggtgaacttcaagatccgccacaacatcgaggacggcagcgtgcagctcgccgaccactacc
agcagaacacccccatcggcgacggccccgtgctgctgcccgacaaccactacctgagcacccagtccgccctgagcaaagaccccaac
gagaagcgcgatcacatggtcctgctggagttcgtgaccgccgccgggatcactctcggcatggacgagctgtacaag (EGFP) –
gcggccgcc (linker) -
atggctactcaagctgatttgatggagttggacatggccatggaaccagacagaaaagcggctgttagtcactggcagcaacagtcttacctgga
ctctggaatccattctggtgccactaccacagctccttctctgagtggtaaaggcaatcctgaggaagaggatgtggatacctcccaagtcctgt
atgagtgggaacagggattttctcagtccttcactcaagaacaagtagctgatattgatggacagtatgcaatgactcgagctcagagggtacgag
ctgctatgttccctgagacattagatgagggcatgcagatcccatctacacagtttgatgctgctcatcccactaatgtccagcgtttggctgaac
catcacagatgctgaaacatgcagttgtaaacttgattaactatcaagatgatgcagaacttgccacacgtgcaatccctgaactgacaaaact
gctaaatgacgaggaccaggtggtggttaataaggctgcagttatggtccatcagctttctaaaaaggaagcttccagacacgctatcatgcgttc
tcctcagatggtgtctgctattgtacgtaccatgcagaatacaaatgatgtagaaacagctcgttgtaccgctgggaccttgcataacctttcccat
catcgtgagggcttactggccatctttaagtctggaggcattcctgccctggtgaaaatgcttggttcaccagtggattctgtgttgttttatgccatta
caactctccacaaccttttattacatcaagaaggagctaaaatggcagtgcgtttagctggtgggctgcagaaaatggttgccttgctcaacaaa
acaaatgttaaattcttggctattacgacagactgccttcaaattttagcttatggcaaccaagaaagcaagctcatcatactggctagtggtgga
ccccaagctttagtaaatataatgaggacctatacttacgaaaaactactgtggaccacaagcagagtgctgaaggtgctatctgtctgctctagt
aataagccggctattgtagaagctggtggaatgcaagctttaggacttcacctgacagatccaagtcaacgtcttgttcagaactgtctttggactc
tcaggaatctttcagatgctgcaactaaacaggaagggatggaaggtctccttgggactcttgttcagcttctgggttcagatgatataaatgtggtc
acctgtgcagctggaattctttctaacctcacttgcaataattataagaacaagatgatggtctgccaagtgggtggtatagaggctcttgtgcgta
ctgtccttcgggctggtgacagggaagacatcactgagcctgccatctgtgctcttcgtcatctgaccagccgacaccaagaagcagagatggc
ccagaatgcagttcgccttcactatggactaccagttgtggttaagctcttacacccaccatcccactggcctctgataaaggctactgttggatt
gattcgaaatcttgccctttgtcccgcaaatcatgcacctttgcgtgagcagggtgccattccacgactagttcagttgcttgttcgtgcacatcag
gatacccagcgccgtacgtccatgggtgggacacagcagcaatttgtggagggggtccgcatggaagaaatagttgaaggttgtaccggagccc
ttcacatcctagctcgggatgttcacaaccgaattgttatcagaggactaaataccattccattgtttgtgcagctgctttattctcccattgaaaac
atccaaagagtagctgcaggggtcctctgtgaacttgctcaggacaaggaagctgcagaagctattgaagctgagggagccacagctcctctga
cagagttacttcactctaggaatggaggtgtggcgacatatgcagctgctgttttgttccgaatgtctgaggacaagccacaagattacaagaaac
ggctttcagttgagctgaccagctctctcttcagaacagagccaatggcttggaatgagactgctgatcttggacttgatattggtgcccagggaga
accccttggatatcgccaggatgatcctagctatcgttcttttcactctggtggatatggccaggatgccttgggtatggaccccatgatggaacat
gagatgggtggccaccaccctggtgctgactatccagttgatgggctgccagatctggggcatgcccaggacctcatggatgggctgcctccag
gtgacagcaatcagctggcctggtttgatactgacctgtag (CTNNB1 wild type)

**CTNNB1 (wild type)-GFP**
atggctactcaagctgatttgatggagttggacatggccatggaaccagacagaaaagcggctgttagtcactggcagcaacagtcttacctgga
ctctggaatccattctggtgccactaccacagctccttctctgagtggtaaaggcaatcctgaggaagaggatgtggatacctcccaagtcctgt
atgagtgggaacagggattttctcagtccttcactcaagaacaagtagctgatattgatggacagtatgcaatgactcgagctcagagggtacgag
ctgctatgttccctgagacattagatgagggcatgcagatcccatctacacagtttgatgctgctcatcccactaatgtccagcgtttggctgaac
catcacagatgctgaaacatgcagttgtaaacttgattaactatcaagatgatgcagaacttgccacacgtgcaatccctgaactgacaaaact
gctaaatgacgaggaccaggtggtggttaataaggctgcagttatggtccatcagctttctaaaaaggaagcttccagacacgctatcatgcgttc
tcctcagatggtgtctgctattgtacgtaccatgcagaatacaaatgatgtagaaacagctcgttgtaccgctgggaccttgcataacctttcccat
catcgtgagggcttactggccatctttaagtctggaggcattcctgccctggtgaaaatgcttggttcaccagtggattctgtgttgttttatgccatta
caactctccacaaccttttattacatcaagaaggagctaaaatggcagtgcgtttagctggtgggctgcagaaaatggttgccttgctcaacaaa
acaaatgttaaattcttggctattacgacagactgccttcaaattttagcttatggcaaccaagaaagcaagctcatcatactggctagtggtgga
ccccaagctttagtaaatataatgaggacctatacttacgaaaaactactgtggaccacaagcagagtgctgaaggtgctatctgtctgctctagt
aataagccggctattgtagaagctggtggaatgcaagctttaggacttcacctgacagatccaagtcaacgtcttgttcagaactgtctttggactc
tcaggaatctttcagatgctgcaactaaacaggaagggatggaaggtctccttgggactcttgttcagcttctgggttcagatgatataaatgtggtc
acctgtgcagctggaattctttctaacctcacttgcaataattataagaacaagatgatggtctgccaagtgggtggtatagaggctcttgtgcgta

ctgtccttcgggctggtgacagggaagacatcactgagcctgccatctgtgctcttcgtcatctgaccagccgacaccaagaagcagagatggc
ccagaatgcagttcgccttcactatggactaccagttgtggttaagctcttacacccaccatcccactggcctctgataaaggctactgttggatt
gattcgaaatcttgccctttgtcccgcaaatcatgcacctttgcgtgagcagggtgccattccacgactagttcagttgcttgttcgtgcacatcag
gatacccagcgccgtacgtccatgggtgggacacagcagcaatttgtggagggggtccgcatggaagaaatagttgaaggttgtaccggagccc
ttcacatcctagctcgggatgttcacaaccgaattgttatcagaggactaaataccattccattgtttgtgcagctgctttattctcccattgaaaac
atccaaagagtagctgcaggggtcctctgtgaacttgctcaggacaaggaagctgcagaagctattgaagctgagggagccacagctcctctga
cagagttacttcactctaggaatggaggtgtggcgacatatgcagctgctgttttgttccgaatgtctgaggacaagccacaagattacaagaaac
ggctttcagttgagctgaccagctctctcttcagaacagagccaatggcttggaatgagactgctgatcttggacttgatattggtgcccagggaga
accccttggatatcgccaggatgatcctagctatcgttcttttcactctggtggatatggccaggatgccttgggtatggaccccatgatggaacat
gagatgggtggccaccaccctggtgctgactatccagttgatgggctgccagatctggggcatgcccaggacctcatggatgggctgcctccag
gtgacagcaatcagctggcctggtttgatactgacctg (CTNNB1 wild type) – ggatcc (linker) –
atggtgagcaagggcgaggagctgttcaccggggtggtgcccatcctggtcgagctggacggcgacgtaaacggccacaagttcagcgtgtccg
gcgagggcgagggcgatgccacctacggcaagctgaccctgaagttcatctgcaccaccggcaagctgcccgtgccctggcccaccctcgtg
accaccctgacctacggcgtgcagtgcttcagccgctaccccgaccacatgaagcagcacgacttcttcaagtccgccatgcccgaaggcta
cgtccaggagcgcaccatcttcttcaaggacgacggcaactacaagacccgcgccgaggtgaagttcgagggcgacaccctggtgaaccgc
atcgagctgaagggcatcgacttcaaggaggacggcaacatcctggggcacaagctggagtacaactacaacagccacaacgtctatatcat
ggccgacaagcagaagaacggcatcaaggtgaacttcaagatccgccacaacatcgaggacggcagcgtgcagctcgccgaccactacca
gcagaacacccccatcggcgacggccccgtgctgctgcccgacaaccactacctgagcacccagtccgccctgagcaaagaccccaacg
agaagcgcgatcacatggtcctgctggagttcgtgaccgccgccgggatcactctcggcatggacgagctgtacaagtaa (EGFP)

**Hibit-ALP5-CTNNB1**
Atggtgagcggctggcggctgttcaagaagattagc (Hibit) – gggagctccggtggctcg (linker) –
ccttctggtcgactcgaggaggaattgaggagacgactttctcca (ALP5) –
agtggcggccgctcgagtggctcgggctcgacctcgggctcgggcacaagtttgtacaaaaaagttggcatg (linker) -
atggctactcaagctgatttgatggagttggacatggccatggaaccagacagaaaagcggctgttagtcactggcagcaacagtcttacctgga
ctctggaatccattctggtgccactaccacagctccttctctgagtggtaaaggcaatcctgaggaagaggatgtggatacctcccaagtcctgt
atgagtgggaacagggattttctcagtccttcactcaagaacaagtagctgatattgatggacagtatgcaatgactcgagctcagagggtacgag
ctgctatgttccctgagacattagatgagggcatgcagatcccatctacacagtttgatgctgctcatcccactaatgtccagcgtttggctgaac
catcacagatgctgaaacatgcagttgtaaacttgattaactatcaagatgatgcagaacttgccacacgtgcaatccctgaactgacaaaact
gctaaatgacgaggaccaggtggtggttaataaggctgcagttatggtccatcagctttctaaaaaggaagcttccagacacgctatcatgcgttc
tcctcagatggtgtctgctattgtacgtaccatgcagaatacaaatgatgtagaaacagctcgttgtaccgctgggaccttgcataacctttcccat
catcgtgagggcttactggccatctttaagtctggaggcattcctgccctggtgaaaatgcttggttcaccagtggattctgtgttgttttatgccatta
caactctccacaaccttttattacatcaagaaggagctaaaatggcagtgcgtttagctggtgggctgcagaaaatggttgccttgctcaacaaa
acaaatgttaaattcttggctattacgacagactgccttcaaattttagcttatggcaaccaagaaagcaagctcatcatactggctagtggtgga
ccccaagctttagtaaatataatgaggacctatacttacgaaaaactactgtggaccacaagcagagtgctgaaggtgctatctgtctgctctagt
aataagccggctattgtagaagctggtggaatgcaagctttaggacttcacctgacagatccaagtcaacgtcttgttcagaactgtctttggactc
tcaggaatctttcagatgctgcaactaaacaggaagggatggaaggtctccttgggactcttgttcagcttctgggttcagatgatataaatgtggtc
acctgtgcagctggaattctttctaacctcacttgcaataattataagaacaagatgatggtctgccaagtgggtggtatagaggctcttgtgcgta
ctgtccttcgggctggtgacagggaagacatcactgagcctgccatctgtgctcttcgtcatctgaccagccgacaccaagaagcagagatggc
ccagaatgcagttcgccttcactatggactaccagttgtggttaagctcttacacccaccatcccactggcctctgataaaggctactgttggatt
gattcgaaatcttgccctttgtcccgcaaatcatgcacctttgcgtgagcagggtgccattccacgactagttcagttgcttgttcgtgcacatcag
gatacccagcgccgtacgtccatgggtgggacacagcagcaatttgtggagggggtccgcatggaagaaatagttgaaggttgtaccggagccc
ttcacatcctagctcgggatgttcacaaccgaattgttatcagaggactaaataccattccattgtttgtgcagctgctttattctcccattgaaaac
atccaaagagtagctgcaggggtcctctgtgaacttgctcaggacaaggaagctgcagaagctattgaagctgagggagccacagctcctctga
cagagttacttcactctaggaatggaggtgtggcgacatatgcagctgctgttttgttccgaatgtctgaggacaagccacaagattacaagaaac
ggctttcagttgagctgaccagctctctcttcagaacagagccaatggcttggaatgagactgctgatcttggacttgatattggtgcccagggaga
accccttggatatcgccaggatgatcctagctatcgttcttttcactctggtggatatggccaggatgccttgggtatggaccccatgatggaacat
gagatgggtggccaccaccctggtgctgactatccagttgatgggctgccagatctggggcatgcccaggacctcatggatgggctgcctccag
gtgacagcaatcagctggCCTGGTTTGATACTGACCTGtag (CTNNB1 wild type)

**GFP-CTNNB1 (S33A)**
Atggtgagcaagggcgaggagctgttcaccggggtggtgcccatcctggtcgagctggacggcgacgtaaacggccacaagttcagcgtgtcc
ggcgagggcgagggcgatgccacctacggcaagctgaccctgaagttcatctgcaccaccggcaagctgcccgtgccctggcccaccctcgt

gaccaccctgacctacggcgtgcagtgcttcagccgctaccccgaccacatgaagcagcacgacttcttcaagtccgccatgcccgaaggct
acgtccaggagcgcaccatcttcttcaaggacgacggcaactacaagacccgcgccgaggtgaagttcgagggcgacaccctggtgaaccg
catcgagctgaagggcatcgacttcaaggaggacggcaacatcctggggcacaagctggagtacaactacaacagccacaacgtctatatca
tggccgacaagcagaagaacggcatcaaggtgaacttcaagatccgccacaacatcgaggacggcagcgtgcagctcgccgaccactacc
agcagaacacccccatcggcgacggccccgtgctgctgcccgacaaccactacctgagcacccagtccgccctgagcaaagaccccaac
gagaagcgcgatcacatggtcctgctggagttcgtgaccgccgccgggatcactctcggcatggacgagctgtacaag (EGFP) -
gcggccgcc (linker)-
atggctactcaagctgatttgatggagttggacatggccatggaaccagacagaaaagcggctgttagtcactggcagcaacagtcttacctgga
c**Gct**ggaatccatTctggtgccactAccacagctcctTctctgagtggtaaaggcaatcctgaggaagaggatgtggatacctcccaagtcctg
tatgagtgggaacagggattttctcagtccttcactcaagaacaagtagctgatattgatggacagtatgcaatgactcgagctcagagggtacga
gctgctatgttccctgagacattagatgagggcatgcagatcccatctacacagtttgatgctgctcatcccactaatgtccagcgtttggctgaac
catcacagatgctgaaacatgcagttgtaaacttgattaactatcaagatgatgcagaacttgccacacgtgcaatccctgaactgacaaaact
gctaaatgacgaggaccaggtggtggttaataaggctgcagttatggtccatcagctttctaaaaaggaagcttccagacacgctatcatgcgttc
tcctcagatggtgtctgctattgtacgtaccatgcagaatacaaatgatgtagaaacagctcgttgtaccgctgggaccttgcataacctttcccat
catcgtgagggcttactggccatctttaagtctggaggcattcctgccctggtgaaaatgcttggttcaccagtggattctgtgttgttttatgccatta
caactctccacaaccttttattacatcaagaaggagctaaaatggcagtgcgtttagctggtgggctgcagaaaatggttgccttgctcaacaaa
acaaatgttaaattcttggctattacgacagactgccttcaaattttagcttatggcaaccaagaaagcaagctcatcatactggctagtggtgga
ccccaagctttagtaaatataatgaggacctatacttacgaaaaactactgtggaccacaagcagagtgctgaaggtgctatctgtctgctctagt
aataagccggctattgtagaagctggtggaatgcaagctttaggacttcacctgacagatccaagtcaacgtcttgttcagaactgtctttggactc
tcaggaatctttcagatgctgcaactaaacaggaagggatggaaggtctccttgggactcttgttcagcttctgggttcagatgatataaatgtggtc
acctgtgcagctggaattctttctaacctcacttgcaataattataagaacaagatgatggtctgccaagtgggtggtatagaggctcttgtgcgta
ctgtccttcgggctggtgacagggaagacatcactgagcctgccatctgtgctcttcgtcatctgaccagccgacaccaagaagcagagatggc
ccagaatgcagttcgccttcactatggactaccagttgtggttaagctcttacacccaccatcccactggcctctgataaaggctactgttggatt
gattcgaaatcttgccctttgtcccgcaaatcatgcacctttgcgtgagcagggtgccattccacgactagttcagttgcttgttcgtgcacatcag
gatacccagcgccgtacgtccatgggtgggacacagcagcaatttgtggagggggtccgcatggaagaaatagttgaaggttgtaccggagccc
ttcacatcctagctcgggatgttcacaaccgaattgttatcagaggactaaataccattccattgtttgtgcagctgctttattctcccattgaaaac
atccaaagagtagctgcaggggtcctctgtgaacttgctcaggacaaggaagctgcagaagctattgaagctgagggagccacagctcctctga
cagagttacttcactctaggaatggaggtgtggcgacatatgcagctgctgttttgttccgaatgtctgaggacaagccacaagattacaagaaac
ggctttcagttgagctgaccagctctctcttcagaacagagccaatggcttggaatgagactgctgatcttggacttgatattggtgcccagggaga
accccttggatatcgccaggatgatcctagctatcgttcttttcactctggtggatatggccaggatgccttgggtatggaccccatgatggaacat
gagatgggtggccaccaccctggtgctgactatccagttgatgggctgccagatctggggcatgcccaggacctcatggatgggctgcctccag
gtgacagcaatcagctggcctggtttgatactgacctgtag (CTNNB1 S33A)

**GFP-CTNNB1 (S37A)**

Atggtgagcaagggcgaggagctgttcaccggggtggtgcccatcctggtcgagctggacggcgacgtaaacggccacaagttcagcgtgtcc
ggcgagggcgagggcgatgccacctacggcaagctgaccctgaagttcatctgcaccaccggcaagctgcccgtgccctggcccaccctcgt
gaccaccctgacctacggcgtgcagtgcttcagccgctaccccgaccacatgaagcagcacgacttcttcaagtccgccatgcccgaaggct
acgtccaggagcgcaccatcttcttcaaggacgacggcaactacaagacccgcgccgaggtgaagttcgagggcgacaccctggtgaaccg
catcgagctgaagggcatcgacttcaaggaggacggcaacatcctggggcacaagctggagtacaactacaacagccacaacgtctatatca
tggccgacaagcagaagaacggcatcaaggtgaacttcaagatccgccacaacatcgaggacggcagcgtgcagctcgccgaccactacc
agcagaacacccccatcggcgacggccccgtgctgctgcccgacaaccactacctgagcacccagtccgccctgagcaaagaccccaac
gagaagcgcgatcacatggtcctgctggagttcgtgaccgccgccgggatcactctcggcatggacgagctgtacaag (EGFP) -
gcggccgcc (linker) -
atggctactcaagctgatttgatggagttggacatggccatggaaccagacagaaaagcggctgttagtcactggcagcaacagtcttacctgga
cTctggaatccat**Gct**ggtgccactAccacagctcctTctctgagtggtaaaggcaatcctgaggaagaggatgtggatacctcccaagtcctg
tatgagtgggaacagggattttctcagtccttcactcaagaacaagtagctgatattgatggacagtatgcaatgactcgagctcagagggtacga
gctgctatgttccctgagacattagatgagggcatgcagatcccatctacacagtttgatgctgctcatcccactaatgtccagcgtttggctgaac
catcacagatgctgaaacatgcagttgtaaacttgattaactatcaagatgatgcagaacttgccacacgtgcaatccctgaactgacaaaact
gctaaatgacgaggaccaggtggtggttaataaggctgcagttatggtccatcagctttctaaaaaggaagcttccagacacgctatcatgcgttc
tcctcagatggtgtctgctattgtacgtaccatgcagaatacaaatgatgtagaaacagctcgttgtaccgctgggaccttgcataacctttcccat
catcgtgagggcttactggccatctttaagtctggaggcattcctgccctggtgaaaatgcttggttcaccagtggattctgtgttgttttatgccatta
caactctccacaaccttttattacatcaagaaggagctaaaatggcagtgcgtttagctggtgggctgcagaaaatggttgccttgctcaacaaa
acaaatgttaaattcttggctattacgacagactgccttcaaattttagcttatggcaaccaagaaagcaagctcatcatactggctagtggtgga

ccccaagctttagtaaatataatgaggacctatacttacgaaaaactactgtggaccacaagcagagtgctgaaggtgctatctgtctgctctagt
aataagccggctattgtagaagctggtggaatgcaagctttaggacttcacctgacagatccaagtcaacgtcttgttcagaactgtctttggactc
tcaggaatctttcagatgctgcaactaaacaggaagggatggaaggtctccttgggactcttgttcagcttctgggttcagatgatataaatgtggtc
acctgtgcagctggaattctttctaacctcacttgcaataattataagaacaagatgatggtctgccaagtgggtggtatagaggctcttgtgcgta
ctgtccttcgggctggtgacagggaagacatcactgagcctgccatctgtgctcttcgtcatctgaccagccgacaccaagaagcagagatggc
ccagaatgcagttcgccttcactatggactaccagttgtggttaagctcttacacccaccatcccactggcctctgataaaggctactgttggatt
gattcgaaatcttgccctttgtcccgcaaatcatgcacctttgcgtgagcagggtgccattccacgactagttcagttgcttgttcgtgcacatcag
gatacccagcgccgtacgtccatgggtgggacacagcagcaatttgtggagggggtccgcatggaagaaatagttgaaggttgtaccggagccc
ttcacatcctagctcgggatgttcacaaccgaattgttatcagaggactaaataccattccattgtttgtgcagctgctttattctcccattgaaaac
atccaaagagtagctgcaggggtcctctgtgaacttgctcaggacaaggaagctgcagaagctattgaagctgagggagccacagctcctctga
cagagttacttcactctaggaatggaggtgtggcgacatatgcagctgctgttttgttccgaatgtctgaggacaagccacaagattacaagaaac
ggctttcagttgagctgaccagctctctcttcagaacagagccaatggcttggaatgagactgctgatcttggacttgatattggtgcccagggaga
accccttggatatcgccaggatgatcctagctatcgttcttttcactctggtggatatggccaggatgccttgggtatggaccccatgatggaacat
gagatgggtggccaccaccctggtgctgactatccagttgatgggctgccagatctggggcatgcccaggacctcatggatgggctgcctccag
gtgacagcaatcagctggcctggtttgatactgacctgtag (CTNNB1 S37A)

**GFP-CTNNB1 (T41A)**

Atggtgagcaagggcgaggagctgttcaccggggtggtgcccatcctggtcgagctggacggcgacgtaaacggccacaagttcagcgtgtcc
ggcgagggcgagggcgatgccacctacggcaagctgaccctgaagttcatctgcaccaccggcaagctgcccgtgccctggcccaccctcgt
gaccaccctgacctacggcgtgcagtgcttcagccgctaccccgaccacatgaagcagcacgacttcttcaagtccgccatgcccgaaggct
acgtccaggagcgcaccatcttcttcaaggacgacggcaactacaagacccgcgccgaggtgaagttcgagggcgacaccctggtgaaccg
catcgagctgaagggcatcgacttcaaggaggacggcaacatcctggggcacaagctggagtacaactacaacagccacaacgtctatatca
tggccgacaagcagaagaacggcatcaaggtgaacttcaagatccgccacaacatcgaggacggcagcgtgcagctcgccgaccactacc
agcagaacacccccatcggcgacggccccgtgctgctgcccgacaaccactacctgagcacccagtccgccctgagcaaagaccccaac
gagaagcgcgatcacatggtcctgctggagttcgtgaccgccgccgggatcactctcggcatggacgagctgtacaag (EGFP) -
gcggccgcc (linker) -
atggctactcaagctgatttgatggagttggacatggccatggaaccagacagaaaagcggctgttagtcactggcagcaacagtcttacctgga
cTctggaatccatTctggtgccact**Gcc**acagctcctTctctgagtggtaaaggcaatcctgaggaagaggatgtggatacctcccaagtcctg
tatgagtgggaacagggattttctcagtccttcactcaagaacaagtagctgatattgatggacagtatgcaatgactcgagctcagagggtacga
gctgctatgttccctgagacattagatgagggcatgcagatcccatctacacagtttgatgctgctcatcccactaatgtccagcgtttggctgaac
catcacagatgctgaaacatgcagttgtaaacttgattaactatcaagatgatgcagaacttgccacacgtgcaatccctgaactgacaaaact
gctaaatgacgaggaccaggtggtggttaataaggctgcagttatggtccatcagctttctaaaaaggaagcttccagacacgctatcatgcgttc
tcctcagatggtgtctgctattgtacgtaccatgcagaatacaaatgatgtagaaacagctcgttgtaccgctgggaccttgcataacctttcccat
catcgtgagggcttactggccatctttaagtctggaggcattcctgccctggtgaaaatgcttggttcaccagtggattctgtgttgttttatgccatta
caactctccacaaccttttattacatcaagaaggagctaaaatggcagtgcgtttagctggtgggctgcagaaaatggttgccttgctcaacaaa
acaaatgttaaattcttggctattacgacagactgccttcaaattttagcttatggcaaccaagaaagcaagctcatcatactggctagtggtgga
ccccaagctttagtaaatataatgaggacctatacttacgaaaaactactgtggaccacaagcagagtgctgaaggtgctatctgtctgctctagt
aataagccggctattgtagaagctggtggaatgcaagctttaggacttcacctgacagatccaagtcaacgtcttgttcagaactgtctttggactc
tcaggaatctttcagatgctgcaactaaacaggaagggatggaaggtctccttgggactcttgttcagcttctgggttcagatgatataaatgtggtc
acctgtgcagctggaattctttctaacctcacttgcaataattataagaacaagatgatggtctgccaagtgggtggtatagaggctcttgtgcgta
ctgtccttcgggctggtgacagggaagacatcactgagcctgccatctgtgctcttcgtcatctgaccagccgacaccaagaagcagagatggc
ccagaatgcagttcgccttcactatggactaccagttgtggttaagctcttacacccaccatcccactggcctctgataaaggctactgttggatt
gattcgaaatcttgccctttgtcccgcaaatcatgcacctttgcgtgagcagggtgccattccacgactagttcagttgcttgttcgtgcacatcag
gatacccagcgccgtacgtccatgggtgggacacagcagcaatttgtggagggggtccgcatggaagaaatagttgaaggttgtaccggagccc
ttcacatcctagctcgggatgttcacaaccgaattgttatcagaggactaaataccattccattgtttgtgcagctgctttattctcccattgaaaac
atccaaagagtagctgcaggggtcctctgtgaacttgctcaggacaaggaagctgcagaagctattgaagctgagggagccacagctcctctga
cagagttacttcactctaggaatggaggtgtggcgacatatgcagctgctgttttgttccgaatgtctgaggacaagccacaagattacaagaaac
ggctttcagttgagctgaccagctctctcttcagaacagagccaatggcttggaatgagactgctgatcttggacttgatattggtgcccagggaga
accccttggatatcgccaggatgatcctagctatcgttcttttcactctggtggatatggccaggatgccttgggtatggaccccatgatggaacat
gagatgggtggccaccaccctggtgctgactatccagttgatgggctgccagatctggggcatgcccaggacctcatggatgggctgcctccag
gtgacagcaatcagctggcctggtttgatactgacctgtag (CTNNB1 T41A)

**GFP-CTNNB1 (S45del, CSNK phosphorylation site, this deletion is found in the colorectal cancer cell line HCT116)**

Atggtgagcaagggcgaggagctgttcaccggggtggtgcccatcctggtcgagctggacggcgacgtaaacggccacaagttcagcgtgtcc
ggcgagggcgagggcgatgccacctacggcaagctgaccctgaagttcatctgcaccaccggcaagctgcccgtgccctggcccaccctcgt
gaccaccctgacctacggcgtgcagtgcttcagccgctaccccgaccacatgaagcagcacgacttcttcaagtccgccatgcccgaaggct
acgtccaggagcgcaccatcttcttcaaggacgacggcaactacaagacccgcgccgaggtgaagttcgagggcgacaccctggtgaaccg
catcgagctgaagggcatcgacttcaaggaggacggcaacatcctggggcacaagctggagtacaactacaacagccacaacgtctatatca
tggccgacaagcagaagaacggcatcaaggtgaacttcaagatccgccacaacatcgaggacggcagcgtgcagctcgccgaccactacc
agcagaacacccccatcggcgacggccccgtgctgctgcccgacaaccactacctgagcacccagtccgccctgagcaaagaccccaac
gagaagcgcgatcacatggtcctgctggagttcgtgaccgccgccgggatcactctcggcatggacgagctgtacaag (EGFP) -
gcggccgcc (linker) -
atggctactcaagctgatttgatggagttggacatggccatggaaccagacagaaaagcggctgttagtcactggcagcaacagtcttacctgga
cTctggaatccatTctggtgccactAccacagctcctctgagtggtaaaggcaatcctgaggaagaggatgtggatacctcccaagtcctgtatg
agtgggaacagggattttctcagtccttcactcaagaacaagtagctgatattgatggacagtatgcaatgactcgagctcagagggtacgagctg
ctatgttccctgagacattagatgagggcatgcagatcccatctacacagtttgatgctgctcatcccactaatgtccagcgtttggctgaaccat
cacagatgctgaaacatgcagttgtaaacttgattaactatcaagatgatgcagaacttgccacacgtgcaatccctgaactgacaaaactgct
aaatgacgaggaccaggtggtggttaataaggctgcagttatggtccatcagctttctaaaaaggaagcttccagacacgctatcatgcgttctcc
tcagatggtgtctgctattgtacgtaccatgcagaatacaaatgatgtagaaacagctcgttgtaccgctgggaccttgcataacctttcccatcat
cgtgagggcttactggccatctttaagtctggaggcattcctgccctggtgaaaatgcttggttcaccagtggattctgtgttgttttatgccattacaa
ctctccacaaccttttattacatcaagaaggagctaaaatggcagtgcgtttagctggtgggctgcagaaaatggttgccttgctcaacaaaaca
aatgttaaattcttggctattacgacagactgccttcaaattttagcttatggcaaccaagaaagcaagctcatcatactggctagtggtggaccc
caagctttagtaaatataatgaggacctatacttacgaaaaactactgtggaccacaagcagagtgctgaaggtgctatctgtctgctctagtaat
aagccggctattgtagaagctggtggaatgcaagctttaggacttcacctgacagatccaagtcaacgtcttgttcagaactgtctttggactctca
ggaatctttcagatgctgcaactaaacaggaagggatggaaggtctccttgggactcttgttcagcttctgggttcagatgatataaatgtggtcac
ctgtgcagctggaattctttctaacctcacttgcaataattataagaacaagatgatggtctgccaagtgggtggtatagaggctcttgtgcgtactg
tccttcgggctggtgacagggaagacatcactgagcctgccatctgtgctcttcgtcatctgaccagccgacaccaagaagcagagatggccc
agaatgcagttcgccttcactatggactaccagttgtggttaagctcttacacccaccatcccactggcctctgataaaggctactgttggattgat
tcgaaatcttgccctttgtcccgcaaatcatgcacctttgcgtgagcagggtgccattccacgactagttcagttgcttgttcgtgcacatcaggat
acccagcgccgtacgtccatgggtgggacacagcagcaatttgtggagggggtccgcatggaagaaatagttgaaggttgtaccggagcccttc
acatcctagctcgggatgttcacaaccgaattgttatcagaggactaaataccattccattgtttgtgcagctgctttattctcccattgaaaacatc
caaagagtagctgcaggggtcctctgtgaacttgctcaggacaaggaagctgcagaagctattgaagctgagggagccacagctcctctgaca
gagttacttcactctaggaatggaggtgtggcgacatatgcagctgctgttttgttccgaatgtctgaggacaagccacaagattacaagaaacgg
ctttcagttgagctgaccagctctctcttcagaacagagccaatggcttggaatgagactgctgatcttggacttgatattggtgcccagggagaac
cccttggatatcgccaggatgatcctagctatcgttcttttcactctggtggatatggccaggatgccttgggtatggaccccatgatggaacatga
gatgggtggccaccaccctggtgctgactatccagttgatgggctgccagatctggggcatgcccaggacctcatggatgggctgcctccaggtg
acagcaatcagctggcctggtttgatactgacctgtag (CTNNB1 S45del)

**GFP-CTNNB1 (S33A/S37A, beta-TrCP binding sites)**

Atggtgagcaagggcgaggagctgttcaccggggtggtgcccatcctggtcgagctggacggcgacgtaaacggccacaagttcagcgtgtcc
ggcgagggcgagggcgatgccacctacggcaagctgaccctgaagttcatctgcaccaccggcaagctgcccgtgccctggcccaccctcgt
gaccaccctgacctacggcgtgcagtgcttcagccgctaccccgaccacatgaagcagcacgacttcttcaagtccgccatgcccgaaggct
acgtccaggagcgcaccatcttcttcaaggacgacggcaactacaagacccgcgccgaggtgaagttcgagggcgacaccctggtgaaccg
catcgagctgaagggcatcgacttcaaggaggacggcaacatcctggggcacaagctggagtacaactacaacagccacaacgtctatatca
tggccgacaagcagaagaacggcatcaaggtgaacttcaagatccgccacaacatcgaggacggcagcgtgcagctcgccgaccactacc
agcagaacacccccatcggcgacggccccgtgctgctgcccgacaaccactacctgagcacccagtccgccctgagcaaagaccccaac
gagaagcgcgatcacatggtcctgctggagttcgtgaccgccgccgggatcactctcggcatggacgagctgtacaag (EGFP) -
gcggccgcc (linker) -
atggctactcaagctgatttgatggagttggacatggccatggaaccagacagaaaagcggctgttagtcactggcagcaacagtcttacctgga
c**Gct**ggaatccat**Gct**ggtgccactAccacagctcctTctctgagtggtaaaggcaatcctgaggaagaggatgtggatacctcccaagtcct
gtatgagtgggaacagggattttctcagtccttcactcaagaacaagtagctgatattgatggacagtatgcaatgactcgagctcagagggtacg
agctgctatgttccctgagacattagatgagggcatgcagatcccatctacacagtttgatgctgctcatcccactaatgtccagcgtttggctgaa
ccatcacagatgctgaaacatgcagttgtaaacttgattaactatcaagatgatgcagaacttgccacacgtgcaatccctgaactgacaaaac
tgctaaatgacgaggaccaggtggtggttaataaggctgcagttatggtccatcagctttctaaaaaggaagcttccagacacgctatcatgcgtt

ctcctcagatggtgtctgctattgtacgtaccatgcagaatacaaatgatgtagaaacagctcgttgtaccgctgggaccttgcataacctttccc
atcatcgtgagggcttactggccatctttaagtctggaggcattcctgccctggtgaaaatgcttggttcaccagtggattctgtgttgttttatgccat
tacaactctccacaaccttttattacatcaagaaggagctaaaatggcagtgcgtttagctggtgggctgcagaaaatggttgccttgctcaacaa
aacaaatgttaaattcttggctattacgacagactgccttcaaattttagcttatggcaaccaagaaagcaagctcatcatactggctagtggtgg
accccaagctttagtaaatataatgaggacctatacttacgaaaaactactgtggaccacaagcagagtgctgaaggtgctatctgtctgctcta
gtaataagccggctattgtagaagctggtggaatgcaagctttaggacttcacctgacagatccaagtcaacgtcttgttcagaactgtctttggac
tctcaggaatctttcagatgctgcaactaaacaggaagggatggaaggtctccttgggactcttgttcagcttctgggttcagatgatataaatgtgg
tcacctgtgcagctggaattctttctaacctcacttgcaataattataagaacaagatgatggtctgccaagtgggtggtatagaggctcttgtgcgt
actgtccttcgggctggtgacagggaagacatcactgagcctgccatctgtgctcttcgtcatctgaccagccgacaccaagaagcagagatgg
cccagaatgcagttcgccttcactatggactaccagttgtggttaagctcttacacccaccatcccactggcctctgataaaggctactgttggat
tgattcgaaatcttgccctttgtcccgcaaatcatgcacctttgcgtgagcagggtgccattccacgactagttcagttgcttgttcgtgcacatcag
gatacccagcgccgtacgtccatgggtgggacacagcagcaatttgtggagggggtccgcatggaagaaatagttgaaggttgtaccggagccc
ttcacatcctagctcgggatgttcacaaccgaattgttatcagaggactaaataccattccattgtttgtgcagctgctttattctcccattgaaaac
atccaaagagtagctgcaggggtcctctgtgaacttgctcaggacaaggaagctgcagaagctattgaagctgagggagccacagctcctctga
cagagttacttcactctaggaatggaggtgtggcgacatatgcagctgctgttttgttccgaatgtctgaggacaagccacaagattacaagaaac
ggctttcagttgagctgaccagctctctcttcagaacagagccaatggcttggaatgagactgctgatcttggacttgatattggtgcccagggaga
accccttggatatcgccaggatgatcctagctatcgttcttttcactctggtggatatggccaggatgccttgggtatggaccccatgatggaacat
gagatgggtggccaccaccctggtgctgactatccagttgatgggctgccagatctggggcatgcccaggacctcatggatgggctgcctccag
gtgacagcaatcagctggcctggtttgatactgacctgtag (CTNNB1 S33A/S37A)

**GFP-CTNNB1 (S33A/S37A/T41A, GSK3 phosphorylation sites)**

Atggtgagcaagggcgaggagctgttcaccggggtggtgcccatcctggtcgagctggacggcgacgtaaacggccacaagttcagcgtgtcc
ggcgagggcgagggcgatgccacctacggcaagctgaccctgaagttcatctgcaccaccggcaagctgcccgtgccctggcccaccctcgt
gaccaccctgacctacggcgtgcagtgcttcagccgctaccccgaccacatgaagcagcacgacttcttcaagtccgccatgcccgaaggct
acgtccaggagcgcaccatcttcttcaaggacgacggcaactacaagacccgcgccgaggtgaagttcgagggcgacaccctggtgaaccg
catcgagctgaagggcatcgacttcaaggaggacggcaacatcctggggcacaagctggagtacaactacaacagccacaacgtctatatca
tggccgacaagcagaagaacggcatcaaggtgaacttcaagatccgccacaacatcgaggacggcagcgtgcagctcgccgaccactacc
agcagaacacccccatcggcgacggccccgtgctgctgcccgacaaccactacctgagcacccagtccgccctgagcaaagaccccaac
gagaagcgcgatcacatggtcctgctggagttcgtgaccgccgccgggatcactctcggcatggacgagctgtacaag (EGFP) -
gcggccgcc (linker) -
atggctactcaagctgatttgatggagttggacatggccatggaaccagacagaaaagcggctgttagtcactggcagcaacagtcttacctgga
c**Gct**ggaatccat**Gct**ggtgccact**Gcc**acagctcctTctctgagtggtaaaggcaatcctgaggaagaggatgtggatacctcccaagtcc
tgtatgagtgggaacagggattttctcagtccttcactcaagaacaagtagctgatattgatggacagtatgcaatgactcgagctcagagggtac
gagctgctatgttccctgagacattagatgagggcatgcagatcccatctacacagtttgatgctgctcatcccactaatgtccagcgtttggctga
accatcacagatgctgaaacatgcagttgtaaacttgattaactatcaagatgatgcagaacttgccacacgtgcaatccctgaactgacaaaa
ctgctaaatgacgaggaccaggtggtggttaataaggctgcagttatggtccatcagctttctaaaaaggaagcttccagacacgctatcatgcgt
tctcctcagatggtgtctgctattgtacgtaccatgcagaatacaaatgatgtagaaacagctcgttgtaccgctgggaccttgcataacctttccc
atcatcgtgagggcttactggccatctttaagtctggaggcattcctgccctggtgaaaatgcttggttcaccagtggattctgtgttgttttatgccat
tacaactctccacaaccttttattacatcaagaaggagctaaaatggcagtgcgtttagctggtgggctgcagaaaatggttgccttgctcaacaa
aacaaatgttaaattcttggctattacgacagactgccttcaaattttagcttatggcaaccaagaaagcaagctcatcatactggctagtggtgg
accccaagctttagtaaatataatgaggacctatacttacgaaaaactactgtggaccacaagcagagtgctgaaggtgctatctgtctgctcta
gtaataagccggctattgtagaagctggtggaatgcaagctttaggacttcacctgacagatccaagtcaacgtcttgttcagaactgtctttggac
tctcaggaatctttcagatgctgcaactaaacaggaagggatggaaggtctccttgggactcttgttcagcttctgggttcagatgatataaatgtgg
tcacctgtgcagctggaattctttctaacctcacttgcaataattataagaacaagatgatggtctgccaagtgggtggtatagaggctcttgtgcgt
actgtccttcgggctggtgacagggaagacatcactgagcctgccatctgtgctcttcgtcatctgaccagccgacaccaagaagcagagatgg
cccagaatgcagttcgccttcactatggactaccagttgtggttaagctcttacacccaccatcccactggcctctgataaaggctactgttggat
tgattcgaaatcttgccctttgtcccgcaaatcatgcacctttgcgtgagcagggtgccattccacgactagttcagttgcttgttcgtgcacatcag
gatacccagcgccgtacgtccatgggtgggacacagcagcaatttgtggagggggtccgcatggaagaaatagttgaaggttgtaccggagccc
ttcacatcctagctcgggatgttcacaaccgaattgttatcagaggactaaataccattccattgtttgtgcagctgctttattctcccattgaaaac
atccaaagagtagctgcaggggtcctctgtgaacttgctcaggacaaggaagctgcagaagctattgaagctgagggagccacagctcctctga
cagagttacttcactctaggaatggaggtgtggcgacatatgcagctgctgttttgttccgaatgtctgaggacaagccacaagattacaagaaac
ggctttcagttgagctgaccagctctctcttcagaacagagccaatggcttggaatgagactgctgatcttggacttgatattggtgcccagggaga
accccttggatatcgccaggatgatcctagctatcgttcttttcactctggtggatatggccaggatgccttgggtatggaccccatgatggaacat

gagatgggtggccaccaccctggtgctgactatccagttgatgggctgccagatctggggcatgcccaggacctcatggatgggctgcctccag
gtgacagcaatcagctggcctggtttgatactgacctgtag (CTNNB1 S33A/S37A/T41A)

**GFP-CTNNB1 (S33A/S37A/T41A/S45A)**

Atggtgagcaagggcgaggagctgttcaccggggtggtgcccatcctggtcgagctggacggcgacgtaaacggccacaagttcagcgtgtcc
ggcgagggcgagggcgatgccacctacggcaagctgaccctgaagttcatctgcaccaccggcaagctgcccgtgccctggcccaccctcgt
gaccaccctgacctacggcgtgcagtgcttcagccgctaccccgaccacatgaagcagcacgacttcttcaagtccgccatgcccgaaggct
acgtccaggagcgcaccatcttcttcaaggacgacggcaactacaagacccgcgccgaggtgaagttcgagggcgacacactggtgaaccg
catcgagctgaagggcatcgacttcaaggaggacggcaacatcctggggcacaagctggagtacaactacaacagccacaacgtctatatca
tggccgacaagcagaagaacggcatcaaggtgaacttcaagatccgccacaacatcgaggacggcagcgtgcagctcgccgaccactacc
agcagaacacccccatcggcgacggccccgtgctgctgcccgacaaccactacctgagcacccagtccgccctgagcaaagaccccaac
gagaagcgcgatcacatggtcctgctggagttcgtgaccgccgccgggatcactctcggcatggacgagctgtacaag (EGFP) -
gcggccgcc (linker) -
atggctactcaagctgatttgatggagttggacatggccatggaaccagacagaaaagcggctgttagtcactggcagcaacagtcttacctgga
c**Gct**ggaatccat**Gct**ggtgccact**Gcc**acagctcct**Gct**ctgagtggtaaaggcaatcctgaggaagaggatgtggatacctcccaagtc
ctgtatgagtgggaacagggattttctcagtccttcactcaagaacaagtagctgatattgatggacagtatgcaatgactcgagctcagagggta
cgagctgctatgttccctgagacattagatgagggcatgcagatcccatctacacagtttgatgctgctcatcccactaatgtccagcgtttggctg
aaccatcacagatgctgaaacatgcagttgtaaacttgattaactatcaagatgatgcagaacttgccacacgtgcaatccctgaactgacaaa
actgctaaatgacgaggaccaggtggtggttaataaggctgcagttatggtccatcagctttctaaaaaggaagcttccagacacgctatcatgc
gttctcctcagatggtgtctgctattgtacgtaccatgcagaatacaaatgatgtagaaacagctcgttgtaccgctgggaccttgcataacctttc
ccatcatcgtgagggcttactggccatctttaagtctggaggcattcctgccctggtgaaaatgcttggttcaccagtggattctgtgttgttttatgcc
attacaactctccacaaccttttattacatcaagaaggagctaaaatggcagtgcgtttagctggtgggctgcagaaaatggttgccttgctcaac
aaaacaaatgttaaattcttggctattacgacagactgccttcaaattttagcttatggcaaccaagaaagcaagctcatcatactggctagtggt
ggaccccaagctttagtaaatataatgaggacctatacttacgaaaaactactgtggaccacaagcagagtgctgaaggtgctatctgtctgctc
tagtaataagccggctattgtagaagctggtggaatgcaagctttaggacttcacctgacagatccaagtcaacgtcttgttcagaactgtctttgg
actctcaggaatctttcagatgctgcaactaaacaggaagggatggaaggtctccttgggactcttgttcagcttctgggttcagatgatataaatgt
ggtcacctgtgcagctggaattctttctaacctcacttgcaataattataagaacaagatgatggtctgccaagtgggtggtatagaggctcttgtgc
gtactgtccttcgggctggtgacagggaagacatcactgagcctgccatctgtgctcttcgtcatctgaccagccgacaccaagaagcagagat
ggcccagaatgcagttcgccttcactatggactaccagttgtggttaagctcttacacccaccatcccactggcctctgataaaggctactgttgg
attgattcgaaatcttgccctttgtcccgcaaatcatgcacctttgcgtgagcagggtgccattccacgactagttcagttgcttgttcgtgcacatc
aggatacccagcgccgtacgtccatgggtgggacacagcagcaatttgtggagggggtccgcatggaagaaatagttgaaggttgtaccggagc
ccttcacatcctagctcgggatgttcacaaccgaattgttatcagaggactaaataccattccattgtttgtgcagctgctttattctcccattgaaa
acatccaaagagtagctgcaggggtcctctgtgaacttgctcaggacaaggaagctgcagaagctattgaagctgagggggccacagctcctct
gacagagttacttcactctaggaatggaggtgtggcgacatatgcagctgctgttttgttccgaatgtctgaggacaagccacaagattacaagaa
acggctttcagttgagctgaccagctctctcttcagaacagagccaatggcttggaatgagactgctgatcttggacttgatattggtgcccaggga
gaaccccttggatatcgccaggatgatcctagctatcgttcttttcactctggtggatatggccaggatgccttgggtatggaccccatgatggaac
atgagatgggtggccaccaccctggtgctgactatccagttgatgggctgccagatctggggcatgcccaggacctcatggatgggctgcctcca
ggtgacagcaatcagctggcctggtttgatactgacctgtag (CTNNB1 S33A/S37A/T41A/S45A)
